# Pore formation in complex biological membranes: torn between evolutionary needs

**DOI:** 10.1101/2024.05.06.592649

**Authors:** Leonhard J. Starke, Christoph Allolio, Jochen S. Hub

## Abstract

The primary function of biological membranes is to enable compartmentalization among cells and organelles. Loss of integrity by the formation of membrane pores would trigger uncontrolled depolarization or influx of toxic compounds, posing a fatal thread to living cells. How the lipid complexity of biological membranes enables mechanical stability against pore formation while simultaneously allowing ongoing membrane remodeling is largely enigmatic. We performed molecular dynamics simulations of eight complex lipid membranes including the plasma membrane and membranes of the organelles ER, Golgi, lysosome, and mitochondrion. To quantify the mechanical stability of these membranes, we computed the free energies for nucleating a transmembrane pore as well as the line tension along the rim of open pores. Our simulations reveal that complex biological membranes are overall remarkably stable, however with the plasma membrane standing out as exceptionally stable, which aligns with its crucial role as a protective layer. We observe that sterol content is the main regulator for biomembrane stability, and that lateral sorting among lipid mixtures influences the energetics of membrane pores. A comparison of 25 model membranes with varying sterol content, tail length, tail saturation, and head group type shows that the pore nucleation free energy is mostly associated with the lipid tilt modulus, whereas the line tension along the pore rim is determined by the lipid intrinsic curvature. Together, our study provides an atomistic and energetic view on the role of lipid complexity on biomembrane stability.

**Significance statement:** Biomembranes have evolved to fulfill seemingly conflicting requirements. Membranes form a protective layer against bacterial or viral infection and against external mechanical and toxic stress, thus requiring mechanical stability. However, membranes are furthermore involved in ongoing remodeling for homeostasis, signaling, trafficking, and morphogenesis, necessitating a high degree of plasticity. How the chemical diversity of membranes, comprising hundreds of lipid species, contributes to enable both stability and plasticity is not well understood. We used molecular simulations and free energy calculations of pore formation in complex biomembranes to reveal how mechanical and geometric properties of lipids as well as lateral lipid sorting control the integrity of complex membranes.

## Introduction

Biological membranes contain hundreds of distinct lipid species, ^1^ which vary in their head group composition as well as in the length, unsaturation degree, and number of acyl chains. ^2^ In addition to glycerophospholipids, which are the most abundant lipids in eukaryotes, ^2^ membranes contain lipid classes such as sphingolipids, sterols, or single-tailed lipids. Lipid compositions vary not only between different species and cell types but furthermore between cellular organelles. For instance, the sterol content of eukaryotic membranes increases along the secretory pathway and is maximized in the plasma membrane.^2^ The complexity of many membranes is furthermore increased by asymmetric distributions of lipids between the two membrane leaflets.^3,4^ The biological functions and advantages of this chemical diversity are still largely enigmatic.

Biological membranes have evolved to fulfill seemingly conflicting mechanical requirements. On the one hand, membranes are subject to ongoing remodeling during cell division, endo- and exocytosis, or intracellular trafficking, thus requiring a high degree of structural plasticity. On the other hand, biological membranes serve as barriers between cells and their environment and, in eukaryotic cells, between organelles and the cytosol, thus requiring mechanical stability. Such stability is crucial to withstand the formation of undesired membrane pores, which would lead to potentially fatal transmembrane flux of polar solutes and loss of homeostasis. To realize how lipid complexity contributes to reconcile such seemingly conflicting needs —plasticity and stability— it is mandatory to understand how lipid composition and spatial distribution control the free energy landscape of membrane remodeling.

To quantify membrane stability, we here focus on the resistance of membranes against transmembrane pore formation. Membrane pores may form under various biological or experimental conditions. Pores are formed by proteins such as bacterial toxins,^5^ antimicrobial peptides,^6^ or by proteins of the apoptotic machinery.^7^ Pores may be induced mechanically through high osmotic pressure^8^ or by extensive tension in mechanically active tissue. ^9^ During electroporation, pores are formed by external electric fields, as widely used in medicine and biotechnology to deliver bioactive substances into living cells or to kill malignant cells that are not accessible to surgery.^10,11^ Recently it was found that binding of divalent cations to anionic lipids may impose transmembrane potentials and, thereby, trigger pore formation and transmembrane ionic flux.^12,13^

Direct experimental observation of protein-free pores has been limited to large, micron-sized pores^8^ while nanoscopic pores have been detected indirectly by measuring the permeation of ionic or photoactive compounds. ^14^ Thus, molecular dynamics (MD) simulations have been used extensively to obtain structural and energetic insight into nanoscopic pores. ^15^ According to MD simulations, pore nucleation evolves from local membrane thinning via the formation of a thin membrane-spanning water wire towards the formation of a toroidally-shaped, nanometer-sized, hydrophilic transmembrane pore, characterized by tilted lipid arrangements along the pore rim.^15–17^ In absence of external stressors, pore formation involves considerable energetic costs owing to the initial water penetration into the hydrophobic membrane core as well as owing to membrane bending and lipid tilting deformations along the pore rim.^15,16,18–21^ Previous experimental and computational studies highlighted the effects of membrane thickness^22,23^ and cholesterol content^24,25^ on increased membrane stability against pore formation. On the contrary, the addition of anionic 16:0-18:1-phosphatidylglycerol (POPG) into 16:0-18:1-phospatidylcholine (POPC) membranes has been shown to reduce resistance against pore formation. ^26^

Previous MD studies of pore formation mostly focused on single-component membranes^23,27–32^ or on membranes with at most three to four molecular species. ^33^ Only recently, coarse-grained simulations of electroporation in complex plasma membrane models were reported. In these simulations, pores formed predominantly in regions with enriched density of unsaturated lipid tails. ^34^ However, since atomic details such as the hydrogen bond network of transmembrane water wires likely influence the free energy landscape of pore formation, all-atom simulations are required to obtain a quantitative understanding of pore formation in complex membranes.

Here, we used all-atom MD simulations to compute the free energy landscape of pore formation in complex models of eight different biological membranes including an asymmetric plasma membrane model,^35^ models of several cellular organelles^36,37^ and a model composed of *E. coli* lipid extract.^38^ We computed potentials of mean force (PMFs) of pore nucleation and pore expansion to obtain two principal energetic parameters that quantify the membrane stability against defect formation: the pore nucleation free energy and the line tension along the pore rim.^39,40^ The PMFs reveal that the plasma membrane –in particular its outer leaflet– is by far more stable against pore formation compared to membranes of intracellular compartments. To rationalize the wide range of membrane stabilities and to clarify how lipid species, lipid properties, and membrane elastic properties influence pore formation, we computed additional PMFs of pore formation for a large set of one- or two-component model membranes as well as membrane deformation energies in the context of the Helfrich-Kozlov-Hamm^41,42^ continuum theory. We find that pore nucleation is primarily associated with lipid tilting energies, whereas the line tension along the pore is controlled by bending energies. Together, our calculations provide a comprehensive view on energetics, structures, and mechanisms of pore formation in biological and model membranes.

## Results

We selected a set of eight membrane systems with complex lipid composition, which together represent a wide range of biologically relevant membranes. The complex model membranes include asymmetric models of a plasma membrane,^35^ the mitochondrial outer and inner membranes,^37^ as well as symmetric models with lipid composition from the endosomes, lysosome, endoplasmic reticulum (ER), Golgi apparatus, and *E. coli* lipid extract.^38^ As shown in Fig. 1a, the lipid composition greatly varies among these membranes (see also Tables S1–S8). The *E. coli* lipid extract does not contain PC but is mostly composed of phosphatidylethanolamine (PE), phosphatidylglycerol (PG) and cardiolipin (CL).^43^ Among the eukaryotic membranes, the sterol content increases along the secretory pathway with a low abundance in the ER and Golgi membranes, increased abundance in the lysosome and high abundance up to 40% in the plasma membrane. While cholesterol is symmetrically distributed in the plasma membrane model studied here, other lipids are distributed asymmetrically between the inner and the outer leaflet. Phosphatidylserine (PS) and PE lipids are enriched in the inner plasma membrane leaflet whereas sphingomyelin (SM) is enriched in the outer plasma membrane leaflet (Fig. 1A, two left columns). Likewise, the mitochondrial membrane models are asymmetric, with phosphatidylinositol (PI) being enriched in the inner leaflet of the mitochondrial outer membrane and with CL being enriched in the inner leaflet of the mitochondrial inner membrane, where CL plays a role in stabilizing cristae shaped with negative curvature. ^44^ Additional examples for organelle-specific enrichment of low-abundance lipid species are the presence of phosphatidic acid (PA) and diacylglycerol (DAG) in the ER membrane, rationalized by the function of the ER as the main site of lipid synthesis.^2^ However, the lipid fingerprint of different membranes reflect not only biochemical processes but furthermore the need to withstand different degrees of mechanical and toxic stress, likely rationalizing why the plasma membrane is thicker as compared to organelle membrane, as illustrated by the MD simulation snapshots of Figs. 1b/c.

**Figure 1.**
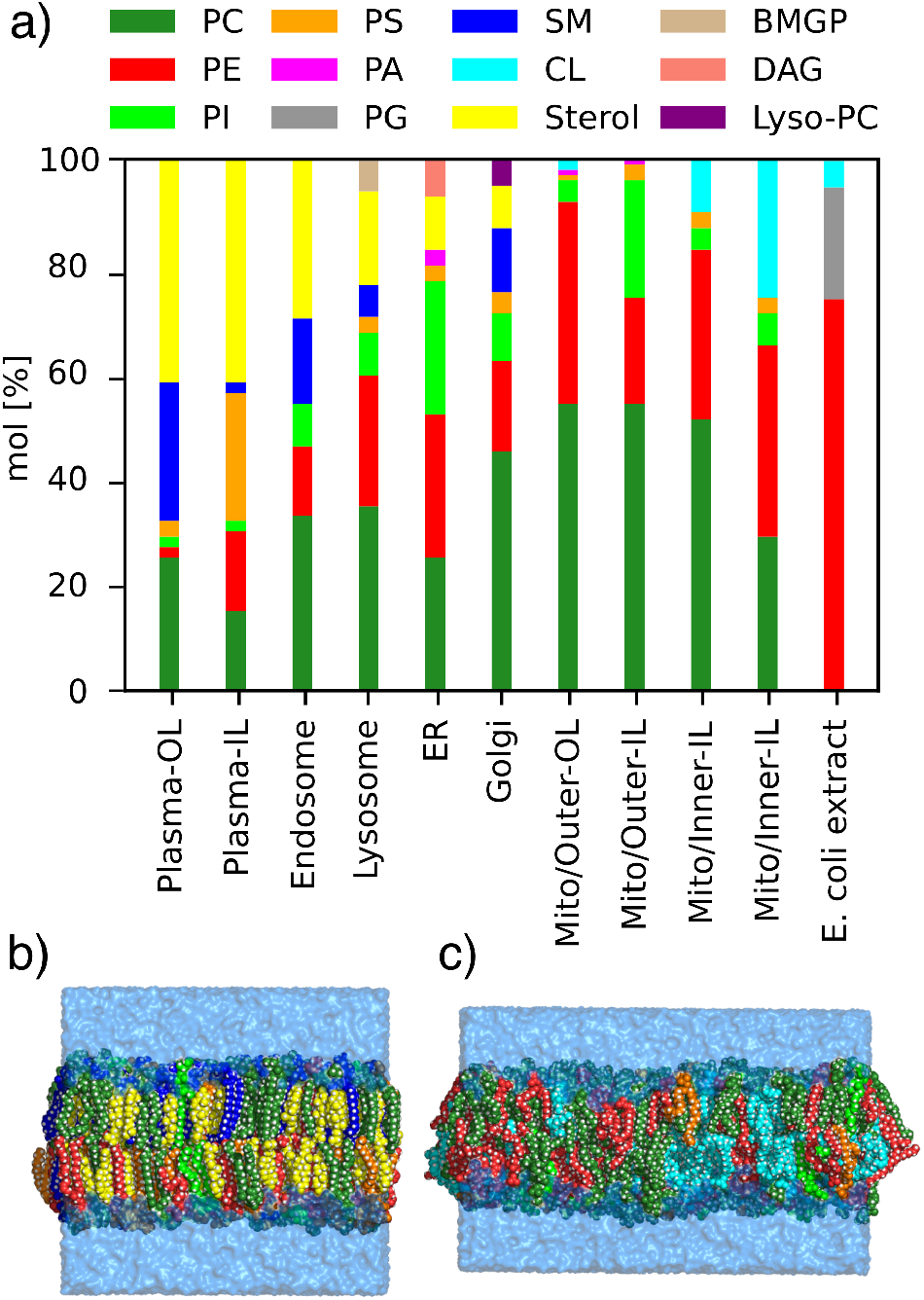
Lipid head group composition of complex membrane systems. (a)Composition of simulated biological membrane systems based on lipid species. PC: phosphatidylcholine, PE: phosphatidylethanolamine, PI: phosphatidylinositol, PS: phosphatidylserine, PA: phosphatidic acid, PG: phosphatidylglycerol, SM: sphingomyelin, CL: cardiolipin, Sterol (cholesterol for mammalian membranes, ergosterol for yeast ER), BMGP: bis(monoacylglycero)phosphate, DAG: diaglycerol, Lyso-PC: lyso-16:0-PC. For asymmetric models of the plasma membrane, mitochondrial outer (Mito/Outer) and inner membranes (Mito/Inner), the composition of the outer leaflet (OL) and the inner leaflet (IL) are shown. (b)Simulation snapshot of the asymmetric plasma membrane and (c) of the asymmetric inner mitochondrial membrane with lipid atoms shown as beads and colored according to the legend in panel (a). Water is represented as transparent light blue surface.

### Pore nucleation free energy Δ*G*_nuc_ and the line tension *γ* along the pore rim

To obtain quantitative and mechanistic insight into pore formation in complex membranes, we computed potentials of mean force (PMFs, also referred to as free energy profiles) of pore formation. PMFs were computed along a joint reaction coordinate *ξ*_p_ of pore nucleation and expansion (see Supplementary Information). ^39,40^ For *ξ*_*p*_ ⪅ 1, the coordinate quantifies the degree of connectivity of a polar transmembrane defect (Fig. 2b–d); ^40^ for *ξ*_*p*_ ⪆ 1, the reaction coordinate quantifies the pore radius *R* in units of the radius *R*_0_ *≈* 0.4 nm of a fully nucleated pore, i.e. *ξ*_p_ = *R/R*_0_ (Fig. 2d–e). Figure 2a presents a typical PMF of pore formation, here shown for the membrane of the Golgi apparatus, plotted either as function of the reaction coordinate *ξ*_p_ (lower abscissa) or, for *ξ*_p_ *>* 1 where a transmembrane defect is present, as function of pore radius *R* (upper abscissa). Starting from the state of a flat membrane (*ξ*_p_ = 0.2, Fig. 2b), the PMF reveals an initial steep increase reflecting the energetic cost for membrane thinning (*ξ*_p_ *≈* 0.6, Fig. 2c). A thin, transmembrane aqueous defect is formed at *ξ*_p_ *≈* 0.9, thereby finalizing the pore nucleation phase (Fig. 2d). The plateau at *ξ*_p_ *≈* 1.5 reflects some energetic stabilization by the tilting of lipids to form a toroidally shaped pore. After further expansion, at *ξ*_p_ ≳ 4 (*R* ≳ 1.5 nm), the PMF increases linearly reflecting mechanical work against an approximately constant line tension *γ* along the pore rim (Fig. 2e). The shape of the PMF and the structural mechanisms involved in pore nucleation and expansion agree qualitatively with results obtained previously for simpler model membranes.^20,39,40^

**Figure 2.**
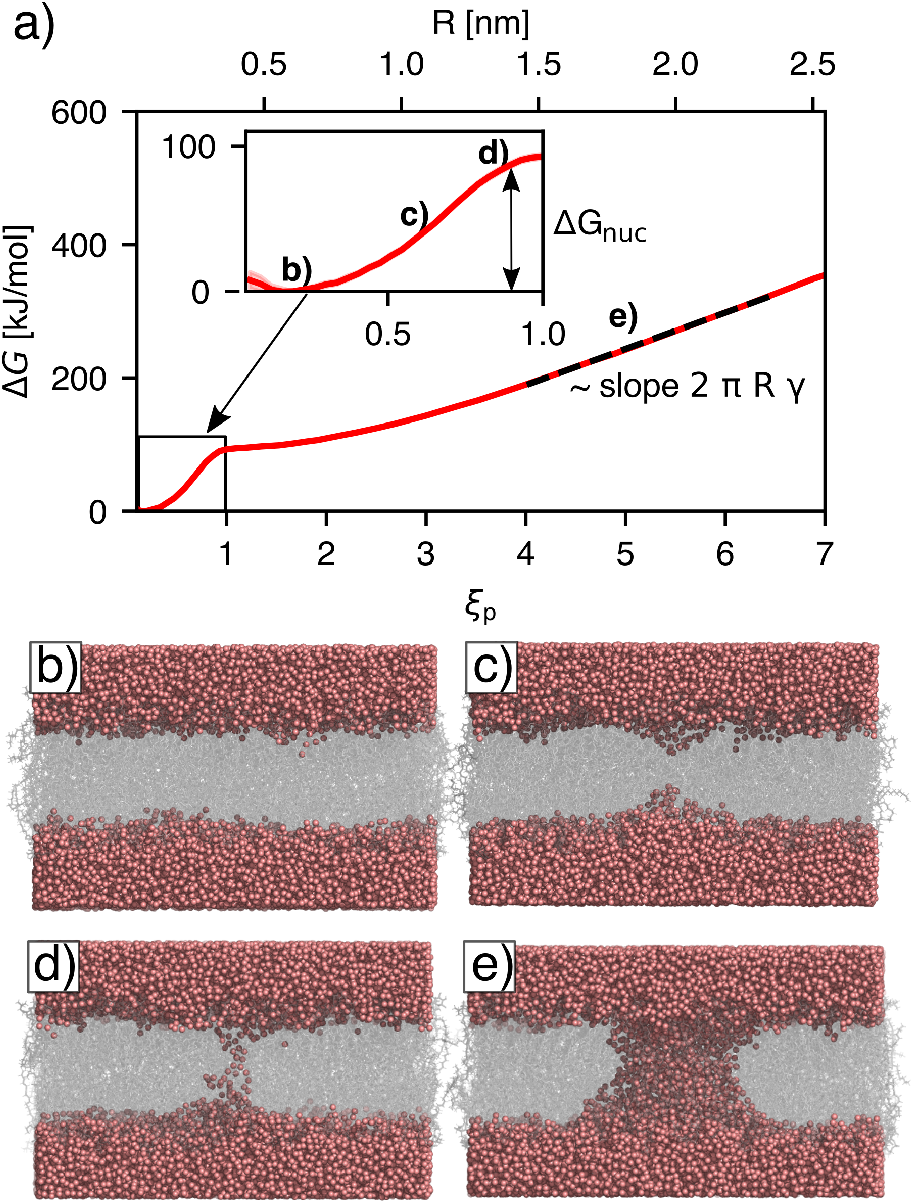
Mechanism and energetics of pore formation. (a)Example PMF of pore nucleation and expansion with the lipid composition of the Golgi apparatus, plotted either as function of the reaction coordinate *ξ*_p_ (lower abscissa) or of pore radius (upper abscissa). The inset shows a closeup view on the PMF during pore nucleation. The nucleation free energy Δ*G*_nuc_ was defined as the PMF value at *ξ*_p_ = 0.92, corresponding to the state with a thin water wire (vertical double arrow). The slope of the linear regime at *ξ*_p_ *>* 4 yields the line tension along the pore rim (black dashed line). Labels b–e indicate structures shown in panels b–e. (b) Simulation snapshot of umbrella windows with: a flat membrane (*ξ*_p_ *≈* 0.2), (c) a thinned membrane (*ξ*_p_ *≈* 0.7), (d) a thin water wire (*ξ*_p_ *≈* 0.9), and (e) a large pore with radius 1.8 nm (*ξ*_p_ *≈* 5). Water oxygen atoms are shown as red spheres, lipids are represented as transparent gray sticks.

The linear regime of the PMF is compatible with classical nucleation theory^45,46^ that describes the free energy of larger pores by Δ*G*_CNT_ = 2*πRγ* − *πR*^2^*σ*, where *γ* denotes the line tension along the pore rim of length 2*πR*. The second term models the relief of surface tension *σ* owing to pore expansion. However, because our simulations were carried out at (i) constant pressure without external surface tension and (ii) at constant number of lipids, thereby keeping the membrane–water interface area approximately constant, the second term can be neglected. Thus, the slope of the linear regime of the PMF at *ξ*_p_ ≳ 4 reflects the line tension *γ* and aligns with classical nucleation theory.

To characterize the energetics of pore nucleation and expansion, we extracted two quantities from the PMFs. Firstly, we defined the pore nucleation energy Δ*G*_nuc_ as the free energy at *ξ*_p_ = 0.92, corresponding by the presence of a thin transmembrane pore (Fig. 2a, inset, black double arrow). Secondly, we obtained the line tension *γ* from the slope of a linear fit to the regime 4 ≤ *ξ*_p_ ≤ 6.5 (Fig. 2a, dashed line). In the following we use Δ*G*_nuc_ and *γ* to compare the free energy landscapes of pore formation of complex or simple model membranes.

### Biological membranes greatly vary in their resistance against pore formation

Among the eight complex biological membranes simulated in this study, the PMFs of pore formation vary considerably (Fig. 3a, colored curves). The nucleation free energy Δ*G*_nuc_ varies between 90 kJ/mol for the Golgi membrane and 180 kJ/mol for the plasma membrane (Fig. 3b). The line tension *γ* varies between 40 pN for the Golgi membrane or for the mitochondrial outer membrane and 96 pN for the plasma membrane (Fig. 3c).

**Figure 3.**
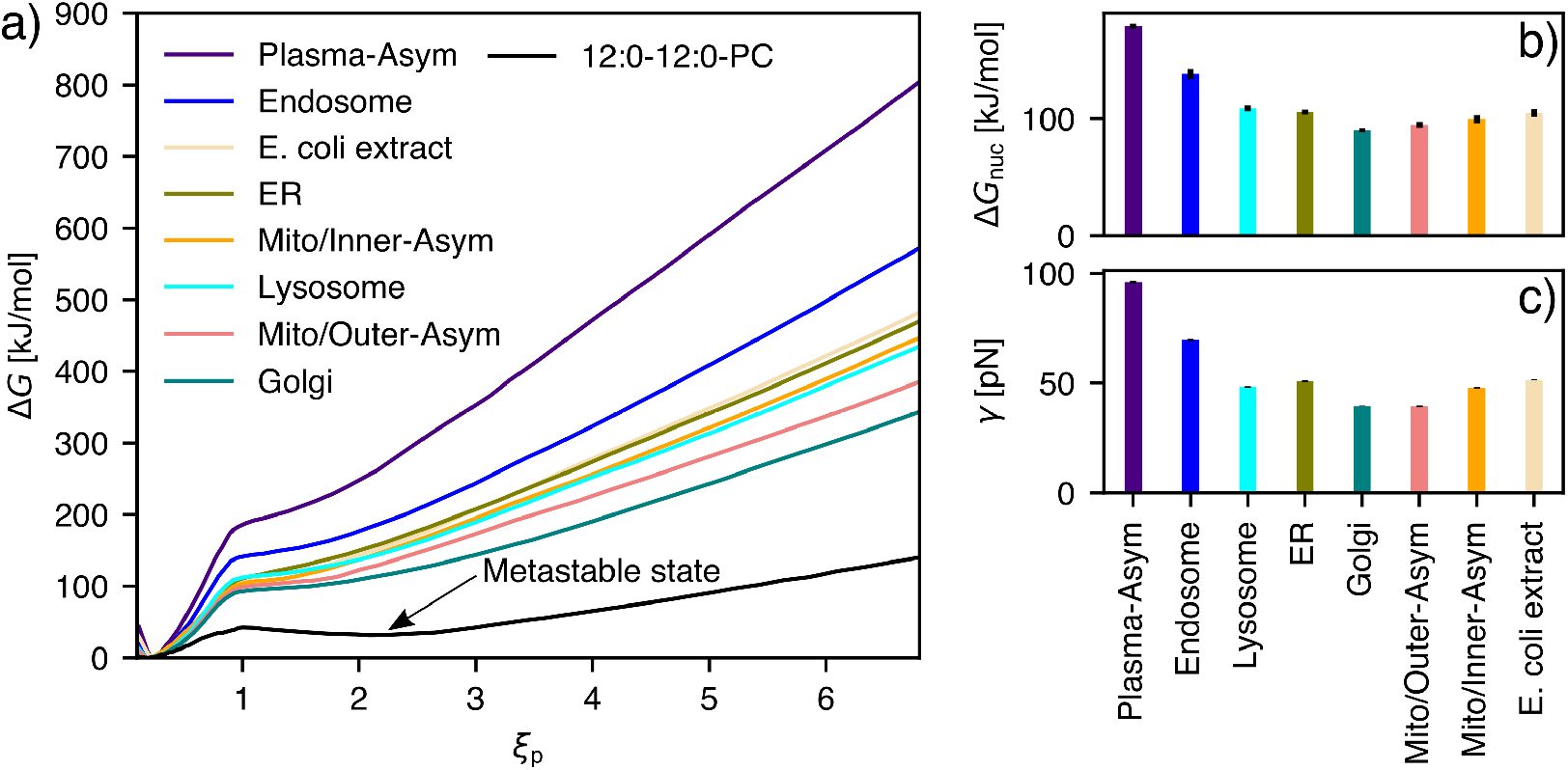
Energetics of pore formation for eight complex membrane systems. (a) PMFs for pore nucleation (*ξ*_p_ *<* 1) and pore expansion (*ξ*_p_ *>* 1) in eight complex biological model membranes and in a membrane of pure 12:0-12:0-PC (DLPC). The arrow highlights a local free energy minimum for the DLPC membrane, indicating a metastable open pore. (b) Pore nucleation free energies Δ*G*_nuc_ and (c) line tension *γ* along the pore rim as taken from the PMFs. Error bars of Δ*G*_nuc_ and *γ* (hardly visible) denote one SE.

These large variations reflect the membranes’ diverse biological functions. Specifically, the plasma membrane is far more stable against pore formation when compared to membranes of intracellular organelles, thus highlighting its ability to form a barrier to the extracellular space. Such marked stability renders the plasma membrane robust against mechanical and osmotic stress, impedes the action of disruptive molecules such as bacterial pore-forming toxins, and forms a barrier against viral infection. In contrast, upwards the secretory pathway, membranes are increasingly unstable against pore formation, as evident from the low Δ*G*_nuc_ and *γ* values of the ER and Golgi membranes as well as, to a lower degree, for the lysosome membrane (Fig. 3b/c). The lower stability of these membranes aligns with their need for increased flexibility to enable ongoing remodeling, while not being exposed to the degree of mechanical or toxic stress that the plasma membrane needs to withstand. Similar arguments rationalize the relatively low stability of the mitochondrial inner and outer membrane. A similarly low stability against pore formation is observed for the membrane formed by *E. coli* extract. However, in the physiological context, Gram-positive and Gram-negative bacteria are surrounded by peptidoglycan and lipopolysaccharide layers, respectively, which strongly contribute to robustness against toxic or mechanical stress.^33,47^ Intermediate stability is observed for early endosomes, which derive from the plasma membrane. ^2^ As shown below, the higher stability of the endosomal membranes is explained by their relatively high cholesterol content of ∼30%, close to the cholesterol content of 40% of the plasma membrane model. Notably, the considerable stability of endosomal membranes may challenge the endosomal escape of drugs or vaccines such as RNA-based therapeutics delivered with lipid nanoparticles.

### Biological membranes do not form metastable pores

Memory effects during early electroporation experiments suggested that lipid membranes may form metastable (long-living) pores^10^ or, equivalently, that pores may represent a local minimum of the free energy landscape. Recently, MD simulations revealed that nanometer-sized pores are metastable only in membranes of inverted cone-shaped lipids. ^32^ For instance, metastable pores are formed by lipids with short saturated tails such as lauroyl acid (di-12:0) or myristoyl acid (di-14:0) and bulky head groups such as PC or PG, whereas unstable pores are formed by lipids with unsaturated tails or smaller head groups form. Whether complex biological membranes from metastable pores has not been systematically addressed.

For reference, Fig. 3a (black curve) presents the PMF of pore formation for di-12:0-PC (DLPC). In agreement with previous studies,^32,39^ the PMF for DLPC reveals a local free energy minimum at *ξ*_p_ *≈* 2.5, implying a metastable pore of radius *R ≈* 1 nm. The presence of the metastable pore may imply pore lifetimes of microseconds or longer.^21,32^ In sharp contrast, no such metastable pore was found in any of the complex membranes, as evident from the absence of corresponding local free energy minimum at *ξ*_p_ *>* 1 (Fig. 3a, colored curves). We hypothesize that the lipid composition of biological membranes has evolved to strictly avoid the formation of long-living pores, underlining that uncontrolled leakage of protons or other ions would be fatal for living cells. Thus, to form long-living pores even in biological membranes, external perturbations are required such as the application of large transmembrane potentials during electroporation experiments^20,27^ or the presence of membrane-active agents including channel-forming toxins,^5^ proteins involved in regulated cell death,^7,48^ polymers,^49^ peptides,^50,51^ or organic solvents such as DMSO. ^39,52^

### The outer leaflet of the plasma membrane is the main protection layer against pore formation

Next, we examined how lipid asymmetry influences pore formation in the plasma membrane and in the inner and outer mitochondrial membranes. To this end, we compared the PMFs obtained with asymmetric membrane models with PMFs obtained with symmetric membranes with the lipid composition of the respective outer or inner leaflet (Fig. S2). The PMFs revealed that the outer plasma membrane leaflet is more stable compared to the inner leaflet, as shown by increased Δ*G*_nuc_ and *γ* values by 42 kJ/mol and 35.7 pN, respectively. These findings suggest that the outer plasma membrane leaflet serves as the primary protection layer against external stressors and against pore-forming agents. However, even the inner leaflet of the plasma membrane is more stable against pore formation when compared to other biological membranes investigated in this study, except for the endosomal membrane that exhibits a similar stability (Fig. S2).

The reduced stability of the inner plasma membrane leaflet aligns with its biological function. Exocytosis involves the formation of a hemifusion stalk between the inner leaflet of the plasma membrane and the outer leaflet of an intracellular vesicle, representing a major structural perturbation of the inner plasma membrane leaflet. Thus, the reduced stability of the inner leaflet against pore formation may provide a compromise between the ability to carry out structural remodeling with acceptable energetic costs while still contributing to the mechanical stability of the overall plasma membrane. These findings are consistent with our previous study that revealed that hemifusion stalk formation by the inner plasma membrane leaflet is far more energetically accessible than stalk formation by the outer leaflet. ^53^

Because sphingomyelin lipids induce increased lipid tail order and membrane stiffness as compared to PC lipids,^54,55^ we hypothesized that sphingomyelin enrichment (Fig. 1a) contributes to the great stability of the plasma membrane outer leaflet. Indeed, by comparing the PMF for a pure POPC membrane with the PMF for a 1:1 mixture of POPC with 18:0-24:0-sphingomyelin (DSM), we found that the addition of DSM increases Δ*G*_nuc_ by 40 kJ/mol (Fig. S3). Thus, in addition to cholesterol whose influence is discussed in the paragraphs below, sphingomyelin contributes to the marked stability of the outer plasma membrane leaflet.

Whereas the two plasma membrane leaflets reveal greatly different stabilities against pore formation, this is not the case for the mitochondrial membranes. Namely, despite the marked asymmetry of the mitochondrial inner and outer membranes (Fig. 1a), symmetric membranes with the composition of the inner or outer leaflet reveal nearly identical Δ*G*_nuc_ and only minor variations of *γ* values (Fig. S2b–e). These findings likely reflect that mitochondrial membranes, relative to the plasma membrane, are faced with far lower mechanical and chemical stress.

### Lipid-specific enrichment and depletion modulate the rim energy

Based on the observation that the addition of a lipid species such as sphingomyelin may strongly alter the free energy of pore formation, we investigated the spatial distribution of lipid species during pore formation. From 400 ns simulations of open pores, we computed the mass densities of several lipid species as function of lateral distance *r* from the pore center and normal distance *z* from the membrane center. Densities were computed for membranes with the composition of the Golgi apparatus (Fig. 4b–i), outer plasma membrane leaflet, endosome, and *E. coli* extract (Fig. S4). The densities reveal enrichment or depletion at the pore rim by several lipid species.

**Figure 4.**
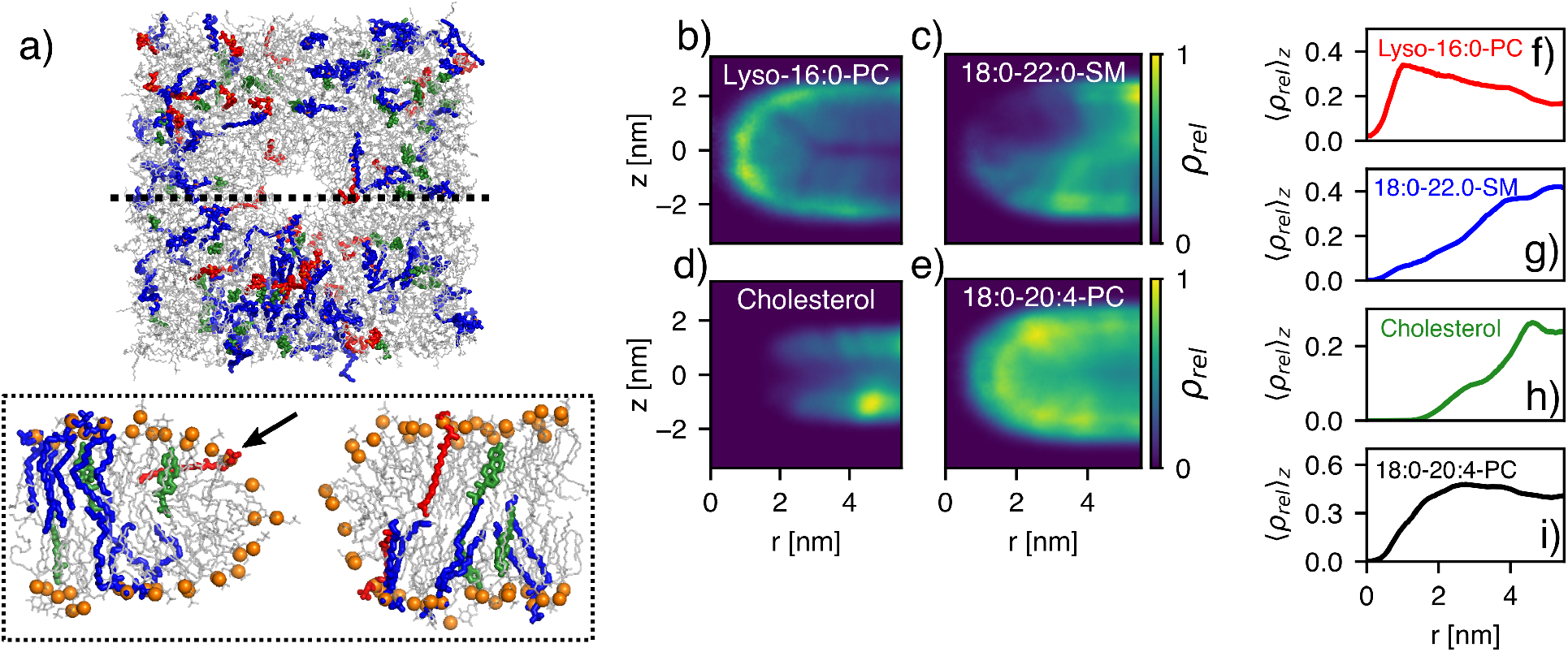
On lipid-specific enrichment or depletion at the pore rim. (a) Simulation snapshot with lipid composition of the Golgi apparatus with a pore with radius *R* = 1.3 nm in top view (upper panel) and side view (lower panel). Lipids are shown as sticks, with phosphorous atoms highlighted as orange spheres. Lyso-16:0-PC is colored in red, cholesterol in green, sphingomyelin in blue, and other lipids in grey. The arrow highlights a tilted lyso-16:0-PC lipid at the pore rim. (b–e) Lipid density along the pore rim as function of radial distance *r* from the pore center and normal distance *z* from the membrane center, relative to the density in the bulk membrane, for (b) lyso-16:0-PC, (c) 18:0-22:0-sphingomyelin, (d) cholesterol, and (e) 18:0-20:4-PC (f-i): One-dimensional lipid density profiles after averaging the densities in panels b–e along *z*.

The toroidal pore rim is characterized by Gaussian curvature with positive curvature along the membrane normal and negative curvature in the membrane plane. Whether the positive or the negative component of the rim curvature controls lipid enrichment and depletion is, *a priori*, not obvious. Among the Golgi membrane lipids, we found that lyso-PC was enriched at the pore rim (Fig. 4b/f), where lyso-PC preferentially adopted a tilted conformation, thereby shielding the hydrophobic membrane core from the transmembrane water defect (Fig. 4a, black arrow). Because lyso-PC is characterized by a large head-to-tail volume ratio, lyso-PC stabilizes positive membrane curvature.^56^ These findings suggest that the positive curvature along the membrane normal dominates the pore energy, and that lipids with intrinsic positive curvature such as lyso-PC may stabilize pores. For instance, lyso-PC likely contributes to the low line tension in the Golgi membrane (Fig. 3c).

In contrast to lyso-PC, cholesterol and sphingomyelin were depleted from pores across the Golgi, E. coli, plasma, and endosomal membranes (Fig. 4a, d/e, g/h and Fig. S4). Thus, the geometries of cholesterol and sphingomyelin are incompatible with the rim structure, which aligns with the large free energies of pore formation in the plasma membrane model that contains 25 % sphingomyelin in the outer leaflet and approx. 40% cholesterol in both leaflets. Thus, enrichment or depletion of specific lipids at the pore rim correlate with stabilizing or destabilizing lipid effects, respectively.

Notably, lateral sorting may lead to different specific lipid concentrations at the pore rim relative to the bulk membrane. Consequently, elastic membrane properties such as bending moduli derived from bulk membranes are not necessarily applicable to the pore rim in complex multi-component membranes.

### Among biological membranes, sterol concentration is the main regulator of membrane stability

Which structural lipid properties control Δ*G*_nuc_ and *γ*, and which lipid properties are employed by biological cells to tune membrane stability? To answer these questions, we investigated systematically how Δ*G*_nuc_ and *γ* are modulated by three structural properties that have been implicated with membrane stability: sterol content,^24,25^ lipid tail length,^22,23^ and degree of unsaturation.^34^ We computed PMFs for twelve model membranes (Fig. S5a–c) and compared the derived Δ*G*_nuc_ and *γ* values (Fig. 5g–l) with the values obtained for complex biological membranes (Fig. 5a–f).

**Figure 5.**
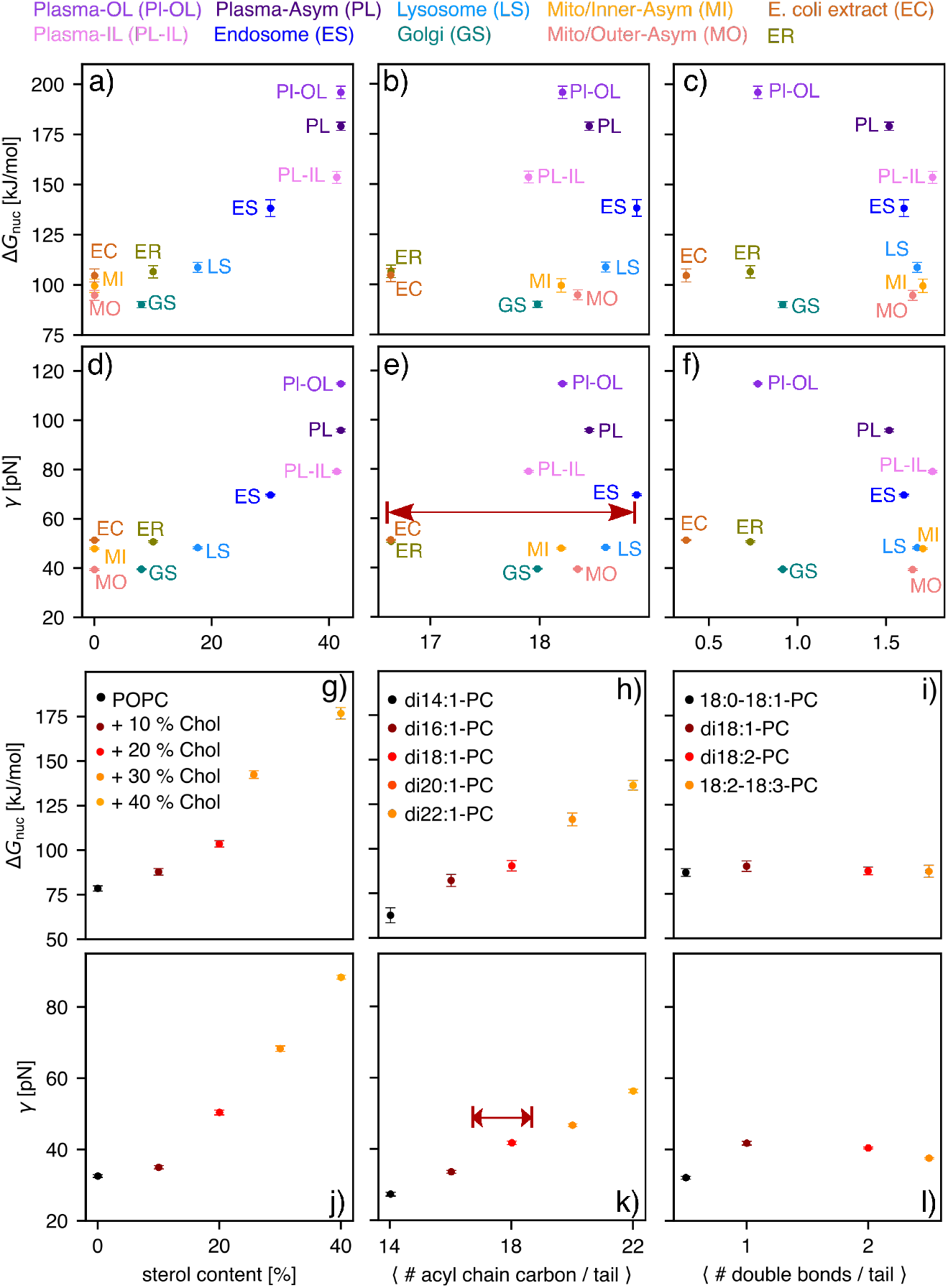
On the effects of sterol concentration (left column), lipid tail length (middle column), and tail unsaturation (right column) on pore nucleation free energy Δ*G*_nuc_ and line tension *γ*. (a-f) Correlation plots (see axis labels) for biological membranes, with abbreviations defined in the upper row. (g–l) Correlation analysis among the same pairs of quantities as shown in panels a–f, however analyzed for model membranes of (g/j) POPC with cholesterol content between 0% and 40%, (h/k) membranes with increasing tail length (see legend), or (i/l) membranes with increasing unsaturation, plotted as number of double bonds per acyl chain (see legend). Dark red arrows highlight the range of average tail lengths realized by biological membranes.

The addition of cholesterol to a membrane of POPC leads a marked concentration-dependent increase of Δ*G*_nuc_ and *γ* by up to 100 kJ/mol and 35 pN, respectively (Fig. 5g/j), which aligns with the trends in complex membranes (Fig. 5a/d). Likewise, by increasing the tail length among five simulations with lipids from di-14:1-PC up to di-22:1-PC, Δ*G*_nuc_ and *γ* increase considerably (Fig. 5h/k), likely owing to increasing membrane thickness.^22,23,57^ Thus, both sterol content and tail length modulate membrane stability against pore formation. However, whereas the cholesterol content varied in the biological membrane models between 0% and 40% (Fig. 5a/d), the average lipid tail length varied in biological membranes by only between 16.7 and 18.8 (Fig. 5e/k, red arrows). These findings demonstrate that biological cells do not make use of the tail length but instead mostly use the sterol content to tune membrane stability.

A recent simulation study based on the coarse-grained MARTINI model reported that poration across complex biological membranes occurs predominantly at sites with a local enrichment of polyunsaturated lipids.^34^ To test the effects of unsaturation on pore formation with atomistic models, we obtained Δ*G*_nuc_ and *γ* for lipids with increasing degree of unsaturation: 18:0-18:1-PC, di-18:1-PC, di-18:2-PC, or 18:2-18:3-PC. We found that the degree of unsaturation has only a small effects on membrane stability against pore formation, thus challenging the interpretation of the coarse-gained simulations. ^34^ While Δ*G*_nuc_ was virtually invariant with respect to unsaturation (Fig. 5i), *γ* revealed an initial rise upon the addition of a second double bond, followed by a minor decrease upon the addition of two or three double bonds per lipid (Fig. 5l). These effects may be rationalized by two opposing effects: on the one hand, a higher degree of unsaturation leads to membrane thinning (Fig. S6b), in line with experimental observations, ^58^ which would lead to decreasing Δ*G*_nuc_ and *γ*. One the other hand, owing to the kinks in the acyl chains, unsaturated lipids induce negative membrane curvature, which is incompatible with the dominating positive curvature along the pore rim discussed above. Thus, effects by polyunsaturated lipids on membrane thickness may cancel with curvature effects, rationalizing the overall low influence on the free energies of pore formation.

### Pore line tension correlates with Helfrich bending energy while nucleation free energy correlates with lipid tilt modulus

Complementary to MD simulations, elasticity theory has been applied to model the free energy of toroidal open pores with contributions from membrane bending, membrane stretching, or lipid tilt.^18,19,41,42,59^ The Helfrich-Kozlov-Hamm (HKH) free energy functional^41,42^ relates the physical properties of lipid monolayers and flat bilayers to the energy of their deformations:

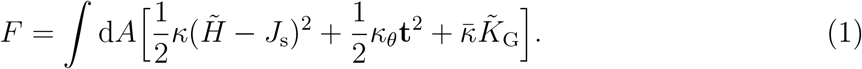

According to the functional, *F* is determined by the bending modulus *κ* that is associated with the difference between the sum 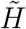 of the two principal curvatures and the spontaneous curvature *J*_s_. The tilde on 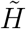 highlights that the calculation is performed over the director field, rather than over the membrane normal vectors. The tilt modulus *κ*_*θ*_ is associated with the local lipid tilt vector **t** defined by the deviation of lipid orientation vector **d** from the membrane normal **n** (Fig. 6c). In the absence of tilt, *F* equals the common Helfrich functional.^41^ The Gaussian curvature 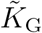 and its associated modulus 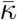 are typically ignored in studies of membrane deformations as, according to the Gauss-Bonnet theorem, the integral over 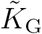 for a closed surface is a topological invariant. However, the formation of a pore either constitutes the formation of a boundary or a new topology, hence this modulus is potentially relevant for pore formation. The HKH theory of membrane elasticity (Eq. 1) can only be expected to provide a semi-quantitative insight into pore formation for several reasons. First, it is not evident whether the pore rim is to be modeled as curved monolayer or as a hole whose boundary is formed by a large number of tilted lipids. Second, Helfrich theory may be taken as a series expansion of the free energy up to second order around a flat reference geometry; thus, whether the theory is applicable to a highly curved pore rim is unclear.

**Figure 6.**
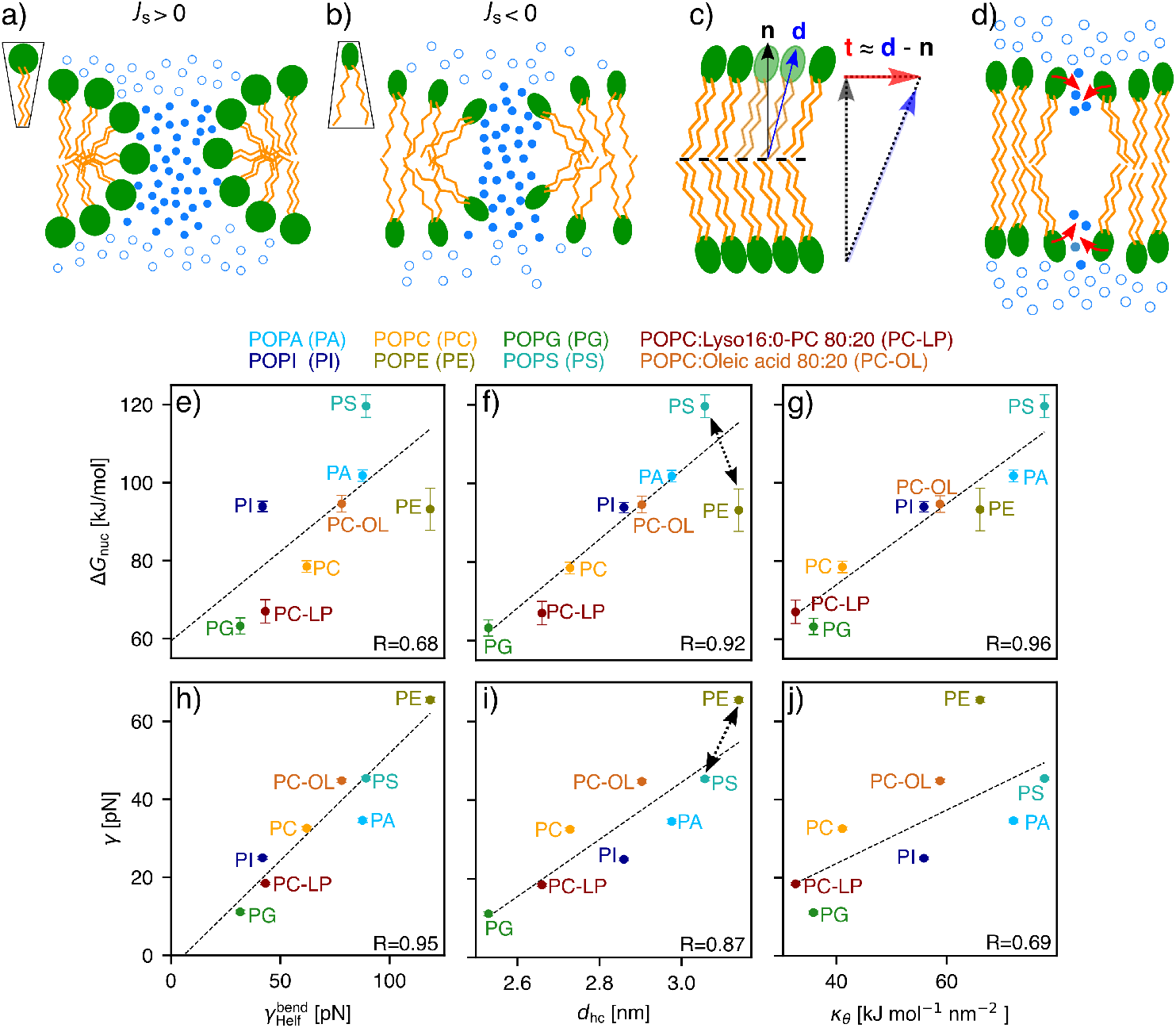
Influence of elastic and geometric properties of the membrane on nucleation free energy Δ*G*_nuc_ and line tension *γ*. (a) Inverted cone-shaped lipids with positive spontaneous curvature *J*_s_ enable favorable packing along the pore rim, whereas (b) cone-shaped lipids with negative *J*_s_ reveal unfavorable packing along the rim. Contribution of bending is characterized by 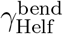. (c) Definition of lipid tilt vector **t** = **d***/*(**n** *·* **d**) − **n** *≈* **d** − **n**, with **n** denoting the membrane normal vector and **d** the director. (d) Illustration of lipid tilting during pore nucleation, with lipid tilting being capable of stabilizing the initial aqueous defect. (e/h, f/i, g/j) Correlations of Δ*G*_nuc_ or *γ* with 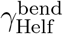, hydrophobic core thickness *d*_hc_ and tilt modulus *κ*_0_ (see axis labels) for six palmitoyl-oleoyl (PO, 16:0-18:1) membranes with varying head groups of PA, PI, PG, PC, PE, or PS, as well as 80:20 mixtures of POPC with 16:0-lyso-PC (PC-LP) or with oleic acid (PC-OL). Results of a linear regression are indicated as dashed lines. Correlation coefficients *R* are shown (see Tab. S10). Dashed arrows (f/i) indicate reversed effect of PS and PE head groups on Δ*G*_nuc_ and *γ*.

Third, for small pores, details of hydrogen bond networks, the finite size of water molecules, and electrostatic interactions may be important.^20^ Nonetheless, the theory has been applied to model pore formation via a variational minimization of the HKH functional with variable pore boundary geometry using a boundary condition in the director orientation. ^19^

In this study, we do not aim to explain pore free energies quantitatively by HKH theory. Instead, we correlate pore free energies from PMF calculations to lipid properties in the context of the HKH theory (such as spontaneous curvature, bending or tilt moduli), with the aim to reveal the physicochemical determinants of membrane stability. To this end, by means of a back-of-the-envelope estimation, membrane bending contributes to the free energy of large pores via the term 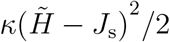 (see Eq. 1), where 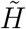 is the large positive curvature along the pore, while *J*_s_ is the spontaneous curvature. The effect of *J*_s_ on pore formation has frequently been interpreted geometrically: ^60^ inverted-cone shaped or cone-shaped lipids impose positive or negative *J*_s_, thereby enabling either favorable or unfavorable packing along the pore rim, respectively (Fig. 6a/b). For instance, unsaturated fatty acids such as oleic acid with a bulky tail and a small polar head group impose negative *J*_s_, whereas saturated mono acyl lipids such as lyso-16:0-PC impose positive *J*_s_.^61^ Thus, oleic acid and lyso-16:0-PC are expected to impede or to facilitate pore formation, respectively, which we confirmed by a series of PMF calculations with membrane with increasing oleic acid or lyso-PC content (Fig. S7). Lipids with *J*_*s*_ *≈* 0 such as POPC are associated with a cylindrical shape, thus expected to show intermediate behavior. Integrating the bending energy functional (first term in Eq. 1) over the rim of a large pore^18^ yields the line tension 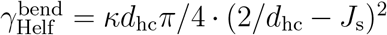, where *d*_hc_ denotes the thickness of the membrane hydrophobic core. Here, it was assumed that one principal curvature is given by *c*_1_ = 1*/*(*d*_hc_*/*2), whereas the second principal curvature is approximately zero for a large pore. Following previous work,^18^ we hypothesized that the line tension *γ* obtained from simulations may be correlated with the line tension 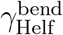 suggested by elasticity theory.

In contrast to wide open pores, the defects during the pore nucleation phase do not adopt a toroidal shape but are instead characterized by thin aqueous protrusions into the hydrophobic membrane core (Fig. 2c).^21,27–29,31,39,62,63^ Visual inspection of the simulations revealed that such aqueous protrusions are stabilized by partial tilting of lipids, thereby reducing the unfavorable contact of the aqueous protrusion with the membrane core (Fig. 6c/d).^17,39^ Thus, we hypothesized that the free energy of pore nucleation is, in addition to membrane thickness,^23,32^ controlled by the energy of lipid tilting as given by the term 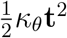 in the HKH functional (Eq. 1).

To systematically investigate the roles of bending and tilting during pore formation, we resorted to simple model membranes of various lipid compositions: single-component model membranes with the same palmitoyl-oleoyl (PO) tails combined with different head groups, namely POPC, POPE, POPS, POPA, and POPG, as well as binary mixtures of POPC with either cholesterol or with the curvature-inducing single-tailed lyso-16:0 PC or oleic acid. From simulations of planar membranes, we computed *J*_s_, *κ*, and *κ*_*θ*_, where values for POPC and POPE were taken from recent studies. ^64,65^ Table 1 demonstrates large variations of *J*_s_ in the range between −0.14 nm^−1^ for POPE, characterizing a cone-shaped lipid,^66^ up to 0.23 nm^−1^ for POPG, characterizing an highly inverted-cone-shaped lipid. As expected, the addition of lyso-16:0-PC or oleic acid to POPC imposes positive or negative *J*_s_, respectively, whereas the effect of cholesterol is hardly detectable within our statistical errors. ^61,67^ Likewise, both the bending modulus *κ* and the tilt modulus *κ*_*θ*_ vary considerably among the model membranes, characterizing harder or softer bending or tilt deformations, respectively. Notably, the addition of 30% cholesterol leads to a strong hardening of both bending and tilt deformations as shown by the by far largest *κ* and *κ*_*θ*_ values. As shown in the following, the variations in *J*_s_, *κ*, and *κ*_*θ*_ have large effects on the free energies of pore formation.

**Table 1:** Spontaneous curvature *J*_s_, monolayer bending modulus *κ*, and bilayer tilt modulus *κ*_*θ*_ computed from planar membranes.

| Component | $J_s$ [$\text{nm}^{-1}$] | $\kappa$ [$\text{kJ mol}^{-1}$] | $\kappa_\theta$ [$\text{kJ mol}^{-1} \text{ nm}^{-2}$] |
| --- | --- | --- | --- |
| POPC<sup>64</sup> | $-0.001 \pm 0.028$ | $33.7 \pm 1.4$ | $42.53 \pm 1.7$ |
| POPE<sup>65</sup> | $-0.14 \pm 0.14$ | $47.7 \pm 2.0$ | $65.6 \pm 2.9$ |
| POPA | $0.019 \pm 0.003$ | $53.1 \pm 2.3$ | $72.1 \pm 2.7$ |
| POPI | $0.187 \pm 0.012$ | $42.8 \pm 2.1$ | $55.8 \pm 2.6$ |
| POPG | $0.23 \pm 0.01$ | $30.2 \pm 1.3$ | $35.8 \pm 1.6$ |
| POPS | $-0.013 \pm 0.001$ | $50.3 \pm 2.3$ | $78 \pm 4$ |
| POPC:lyso16:0-PC 80:20 | $0.087 \pm 0.008$ | $28.2 \pm 1.2$ | $32.6 \pm 1.4$ |
| POPC:oleic acid 80:20 | $-0.051 \pm 0.006$ | $37.7 \pm 1.5$ | $58.8 \pm 2.5$ |
| POPC:Chol 70:30 | $-0.0034 \pm 0.0023$ | $105 \pm 5$ | $166 \pm 8$ |

Figure 6e–j correlates Δ*G*_nuc_ or *γ* obtained from PMFs of pore formation (Fig. S5d/e) with elasticity properties obtained from simulations of planar membranes. In line with our hypothesis stated above, the line tension along the pore rim *γ* strongly correlates with the line tension from HKH theory 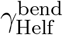 (Fig. 6H, correlation coefficient *R* = 0.95); however, the simple integration of HKH energy functional as carried out here overestimates the line tension by a factor of approximately two, thus emphasizing the need for a consistent minimization including tilt degrees of freedom. The membrane thickness and tilt modulus reveal weaker correlation with *γ* (Fig. 6i/j). This analysis suggests that bending deformation energy is the key contribution to the pore line tension in agreement with previous work. ^19^

In contrast, Δ*G*_nuc_ reveals poor correlation with 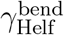 (Fig. 6e) but instead strongly correlates with tilt modulus *κ*_*θ*_ (Fig. 6g, *R* = 0.96). This suggests that soft lipid tilting as found in membranes POPG or POPC:lyso-PC facilitates pore nucleation because head group tilting may stabilize water protrusions into the membrane core, as illustrated in Fig. 6d. Notably, in agreement with previous simulation studies, ^23,32^ Δ*G*_nuc_ furthermore strongly correlates with membrane thickness *d*_hc_. However, the correlation with *d*_hc_ is weaker compared to the correlation with *κ*_*θ*_, mostly because POPE reveals smaller Δ*G*_nuc_ than POPS albeit forming a thicker membrane (Fig. 6f, black arrow). Thus, in addition to membrane thickness, lipid tilting energy emerges as a key determinant for pore nucleation. The importance of thickness and tilting may be expected because they are both proxies for packing of the lipid tails. However, because only the tilt modulus quantifies a cost for modifying the lipid tail packing, the tilt modulus is the better descriptor for the energetic cost of pore formation during which the lipid chain configuration changes dramatically.

## Discussion

We presented PMF calculations of pore formation across complex biological membranes and across simple single- or two-component model membranes, providing two principal energetic parameters: the nucleation free energy Δ*G*_nuc_ and the line tension *γ* along the rim of large pores. The results for *γ* obtained here with the CHARMM36 lipid force field are in reasonable agreement with experiments that reported values between 6.7 and 39.5 pN for various lipid compositions (Table S11).^26,68–72^ However, quantitative comparison is challenging considering that experimental values vary considerably; for instance, reported *γ* values for DOPC vary by up to a factor of four depending on the lipid supplier and reference, which has been attributed to the presence of small amounts of impurities.^68,70^ Nevertheless, trends of *γ* with varying lipid composition observed here agree with previous reports: specifically, *γ* decreased upon replacing PE with PC head groups or by replacing two-tailed PC lipids with lyso-PC.^72^ Likewise, the exceptionally low *γ* and Δ*G*_nuc_ obtained here for POPG aligns with Lira et al., who reported a marked destabilization of giant unilamellar vesicles (GUV) upon the addition of 50% POPG to a membrane of POPC.^26^ Notably, Lira et al. suggested that the low stability of POPG-containing GUVs may indicate general feature of membranes with anionic lipids. In our simulations, in contrast, membrane destabilization was specific for POPG owing to (i) its exceptionally low tilt modulus *κ*_*θ*_, leading to low Δ*G*_nuc_, and (ii) its low bending modulus *κ* together with a large spontaneous curvature *J*_s_, leading to low line tension *γ* (Fig. 6, Table 1 and Table S12). Membranes formed by other anionic lipids POPI, POPA, and POPS exhibited larger Δ*G*_nuc_ and *γ* values, which aligns for the case of POPS with previous experimental and computational studies.^73,74^ However, considering that force fields for anionic lipids have not been refined against experiments as extensively as force fields for zwitterionic lipids, it will be important to revisit the role of anionic lipids during pore formation in future studies.

The nucleation free energies Δ*G*_nuc_ reported above determine the probabilities of pore formation in the small simulation system used during pore nucleation via 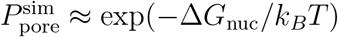, where *k*_*B*_ and *T* denote the Boltzmann constant and temperature, respectively. Because the probability of pore formation is proportional to membrane area, the simulation conditions translate to the free energy of pore formation under experimental conditions of

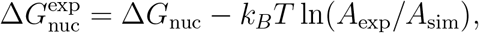

where *A*_exp_ and *A*_sim_ denote the membrane areas in experiment and in our small simulation system, respectively. For instance, for a large GUV of POPG with radius 100 μm, 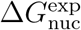 would equal of only 9 kJ/mol, implying frequent water leakage. For the Golgi apparatus with a membrane area of approximately 1000 μm^2^, 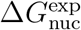 takes a considerable value of 48 kJ/mol, implying a probability of pore formation of 10^−9^ in the absence of any membrane stress.^75^ Thus, considering that the Golgi membrane revealed the smallest Δ*G*_nuc_ value among the biological membranes studied here, these results demonstrate that even biological membranes of inner cellular compartments are remarkably stable, possibly to withstand structural perturbations caused by membrane-associated proteins as well as mechanical stress during membrane remodeling. Notably, the stability of the plasma membrane is increased by another ∼75 kJ/mol relative to the membranes of the inner compartments mostly owing to its high cholesterol and sphingomyelin content, implying another reduction of pore probability by a factor of 10^13^ in the absence of stress. Such marked stability has likely evolved to withstand major mechanical and toxic stress faced by biological cells and, possibly, to reduce the risk of bacterial or viral infection.

We identified cholesterol content as the main regulator for membrane stability among biological membranes. Cholesterol greatly increases *γ* by increasing the bending modulus *κ* (Table 1, Fig. S8) and by moderately increasing the membrane thickness, implying a large bending energy along the pore rim in the context of Helfrich theory. However, cholesterol has only a small effect on the spontaneous curvature *J*_s_ (Tab. 1), suggesting that the membrane-stabilizing effect by cholesterol is not a consequence of intrinsic lipid curvature, but instead is due to an increase in membrane stiffness against both bending and tilting deformations. Hence, concomitant with the the effect on *γ*, cholesterol leads to large Δ*G*_nuc_, in line with an exceptionally large tilt modulus *κ*_*θ*_ (Table 1). Notably, the *γ* and Δ*G*_nuc_ values for the POPC:cholesterol 70:30 membrane are lower than expected from the correlation analysis among other model membranes; we rationalize this finding by lateral depletion of cholesterol from the pore rim (Fig. 4).

## Conclusion

Any form of life relies on tightly controlled compartmentalization by lipid membranes. To quantify the robustness of lipid membranes against mechanical stress, we computed the free energy landscape of pore formation across eight different complex biological membranes, revealing that, both, the nucleation free energy Δ*G*_nuc_ and the line tension along the pore rim *γ* vary greatly among biological membranes. The plasma membrane, and in particular its outer leaflet, is far more stable against pore formation as compared to membranes of intracellular compartments, as shown by Δ*G*_nuc_ and *γ* values increased by ∼75 kJ/mol and ∼45 pN, respectively. However, even membranes of intracellular compartments are remarkably stable against pore formation, quantified by the probability of pore formation per cell in the order of 10^−9^ in the absence of stress, and given that none of the biological membranes studied here form metastable pores. We found that, while membrane stability could in principle be enhanced by increased sterol content or longer lipid tails, biological cells only utilize the sterol content as regulator of membrane stability, likely owing to its simpler biochemical control. Sterol enhances membrane stability via simultaneously increasing membrane thickness, the tilt modulus, and the bending modulus, but not via imposing negative curvature. Compared to sterol content and tail length, lipid polyunsaturation has only a small effect on membrane stability. In complex membranes, the pore free energies may be decreased considerably by enriching or depleting lipids with positive or negative intrinsic curvature at the pore rim, respectively, suggesting that mechanical properties obtained for planar membranes are not applicable to the pore rim region.

To scrutinize the lipid properties that control membrane stability, we computed, for a large set of simpler model membranes, the PMFs of pore formation as well as parameters of membrane elasticity theory, namely the spontaneous curvature, bending modulus, and tilt modulus. We found that *γ* is highly correlated to the bending energy over the pore rim, whereas the biologically important nucleation free energy is primarily determined by the tilt modulus because lipid tilting may stabilize water penetration into the hydrophobic membrane core. Together, our simulations provide an energetic and mechanistic view on transmembrane pore formation, and they reveal how biological cells utilize complex lipid compositions to tune the mechanical stability of membranes, as required for robust compartmentalization.

## Method summary

MD simulations were carried out using GROMACS.^76^ PMF calculations of pore formation were carried out using umbrella sampling (US) along a joint reaction coordinate for pore nucleation and expansion^39,40,77^ implemented in a modified GROMACS version freely available at https://gitlab.com/cbjh/gromacs-chain-coordinate. Lipids were described with the CHARMM36 force field.^78^ Continuum properties, i.e., spontaneous curvature, bending modulus, and tilt modulus were computed from equilibrium simulations of planar membranes as described previously.^64,65,79^ Lipid director definitions as part of the lipidator-toolkit and a modified GROMACS version to calculate the local stress tensor are freely available at https://github.com/allolio. Additional computational details are described in the Supplementary Information, which contains the additional Refs. 80–98.

## Data Availability Statement

A modified GROMACS code that implements the reaction coordinate for pore formation is available at https://gitlab.com/cbjh/gromacs-chain-coordinate. Code for computing mechanical properties of membranes from MD simulations is available at https://github.com/allolio.

## Acknowledgements

C.A. thanks Itay Schachter and Dr. Sukanya Konar for data and discussions. L.S. and J.S.H. were supported by the Deutsche Forschungsgemeinschaft (DFG, German Research Foundation; grants SFB 1027/B7 and INST 256/539-1). C.A. was supported by GAUK PRIMUS Grant PRIMUS/20/SCI/015 and by Charles University Research Centre program No. UNCE/24/SCI/005.

## Notes

### Competing Interest Statement

The authors have declared no competing interest.

