## Supplemental PDF for "Pore formation in complex biological membranes: torn between evolutionary needs"

### Supplementary Methods

#### Setup of membrane systems

Atomistic models of the following complex membranes were set up: (a) asymmetric model of a mammalian plasma membrane,[1] (b) asymmetric models of the outer and (c) inner mitochondrial membranes,[2] symmetric models of (d) the mammalian Golgi apparatus, (e) the lysosome, (f) endosomes,[2] (g) yeast endoplasmic reticulum (ER),[3] and (h) a symmetric model of the *E. coli* polar extract.[4] In addition, to dissect the roles of lipid head groups and tails on the free energy landscape of pore formation, model membranes with simple composition were set up. Single-component membranes were set up containing 16:0-18:1-phosphatidylcholine (POPC), -phosphatidylethanolamine (POPE), -phosphatidylinositol (POPI), -phosphatidylserine (POPS), -phosphatidic acid (POPA), or -phosphatidylglycerol (POPG). Moreover, membranes were set up with binary mixtures of POPC plus cholesterol, 16:0-lyso-PC, oleic acid, or 18:0-24:0-sphingomyelin. For each lipid composition, a small membrane patch with 81 to 162 lipids per leaflet was set up to simulate pore nucleation. Larger membrane patches with 300 to 342 lipids were set up to simulate pore expansion. The lipid compositions of all simulation systems are listed in Tables S1–S8.

Symmetric membranes were generated using MemGen.[5] Asymmetric membrane structures were either taken from CHARMM-GUI[6,7] or built from the symmetric membrane systems. In the latter case the composition of the outer membrane leaflet was adjusted to match the membrane area of the inner leaflet.

Lipid topologies and structures were taken from the CHARMM-GUI web server[6,7] except for *E. coli* polar lipids, which were kindly provided by the authors of Ref. 4.

Atomistic MD simulations were set up and carried out using GROMACS[8] together with the CHARMM36m force field.[9] All systems were neutralized with $Ka^{+}Cl^{-}$ ions and solvated with the CHARMM-modified TIP3P water,[10] which involves Lennard-Jones interactions of hydrogen atoms. The systems were equilibrated for 100 ns. The temperature was controlled at 300 K using the velocity rescale thermostat.[11] The pressure was kept at 1 bar using semi-isotropic exponential pressure relaxation with a stochastic term (c-rescale).[12] Coulomb interactions were calculated using the particle-mesh Ewald method.[13] Lennard-Jones interactions were treated according to the CHARMM specifications, using a cut off at 1.2 nm with the forces being gradually switched off between 1.0 and 1.2 nm. Hydrogen bonds of water molecules were constrained using SETTLE[14] while all other bonds involving hydrogen atoms were constrained using LINCS.[15] Hydrogen mass repartitioning[16] with the default factor of 3 was applied throughout, thus a time step of 4 fs was used for all simulations.

### Reaction coordinate for pore formation

Potentials of mean force (PMF) of pore formation were computed using umbrella sampling (US)[17] along a joint reaction coordinate (RC) $\xi_p$ for pore nucleation and pore expansion.[18] The RC is implemented into an in–house modified version of GROMACS, freely available at `https://gitlab.com/cbjh/gromacs-chain-coordinate`. The RC $\xi_{\mathrm{p}}$ is an extension of the chain coordinate $\xi_{\mathrm{ch}}$ proposed in Ref. 19,20. For graphical illustrations of $\xi_{\mathrm{p}}$ and $\xi_{\mathrm{ch}}$ we refer to Ref. 18. The chain coordinate $\xi_{\mathrm{ch}}$ has been designed to track pore nucleation and quantifies the degree of connectivity of a polar transmembrane defect. $\xi_{\mathrm{ch}}$ is defined using a membrane-spanning narrow cylinder partitioned into $N_{\mathrm{s}}$ slices of thickness $d$. The coordinate evaluates the fraction of slices that are occupied by polar atoms and thus take values in $\xi_{\mathrm{ch}} \in [0, 1)$ with $\xi_{\mathrm{ch}} \approx 1$ indicating a fully formed transmembrane defect. The following parameters were used to specify $\xi_{\mathrm{ch}}$: Both the positions of water oxygen atoms and lipid phosphate moieties were used to compute $\xi_{\mathrm{ch}}$. The radius $R_{\mathrm{cyl}}$ of the cylinder was set to 0.9 nm in this work. The choice of $R_{\mathrm{cyl}}$ ensures that the defect is localized in the membrane plane but does not control the radius of the defect. If $R_{\mathrm{cyl}}$ is too large, two laterally displaced partial defects connected to the upper and lower water reservoirs could be misinterpreted as a single continuous membrane-spanning defect, which could lead to hysteresis problems. The length of the cylinder $N_{\mathrm{s}} \cdot d$ is given by the thickness of the slices (here set to $d = 0.1$ nm) and the number of slices $N_{\mathrm{s}}$. $N_{\mathrm{s}}$ was selected for each membrane such that $\xi_{\mathrm{ch}} \approx 0.25$ was obtained for a flat unperturbed membrane, implying that approximately 25% of the slices (at the head group regions) were filled by polar atoms. The values of $N_{\mathrm{s}}$ used in this study are listed in Table S9.

Because $\xi_{\mathrm{ch}}$ projects larger pores onto $\xi_{\mathrm{ch}} = 1$ it is not suitable for studying pore expansion. Pore expansion is instead characterized by the approximated radius $R$ of the fully formed pore leading to a joint RC $\xi_{\mathrm{p}}$ for pore nucleation and pore expansion,[18] as recently used to obtain the free energy landscape of membrane electroporation.[21,22] The RC $\xi_{\mathrm{p}}$ is defined as follows:

$$\xi_{\mathrm{p}}(\mathbf{r}) = \xi_{\mathrm{ch}}(\mathbf{r}) + H_{\epsilon}\left[\xi_{\mathrm{ch}}(\mathbf{r}) - \xi_{\mathrm{ch}}^{s}\right] \frac{R(\mathbf{r}) - R_0}{R_0}. \tag{1}$$

Here, $\mathbf{r}$ are the Cartesian coordinates of the system. The pore radius $R(\mathbf{r})$ is computed from the polar atoms within a layer of thickness $D$ within the hydrophobic core of the membrane (here set to $D = 1.0$ nm) by assuming a cylindrical shape of the defect and a volume per polar atom of $v_0 = 0.02996\,\mathrm{nm}^3$, corresponding to the molecular volume of water. $R_0$ corresponds to the radius of a minimal transmembrane pore. The value of $R_0$ depends on the membrane system and takes values between 0.32 nm and 0.44 nm (Table S9). $H_\epsilon$ denotes a smoothed differentiable variant of the Heaviside step function, which switches from 0 to 1 within the interval $[-\epsilon, \epsilon]$ (here set to $\epsilon = 0.05$). The value $\xi^s_{\mathrm{ch}}$ determines where to switch from pore nucleation to expansion, here set to $\xi^s_{\mathrm{ch}} = 0.925$. Hence, for $\xi_{\mathrm{ch}} > \xi^s_{\mathrm{ch}} + \epsilon$ corresponding to a fully formed transmembrane defect, we have $H_\epsilon = 1$ and $\xi_{\mathrm{ch}} \approx 1$ and, according to Eq. 1, $\xi_{\mathrm{p}}$ is given by the radius $R$ of the open pore in units of $R_0$ (i.e., $\xi_{\mathrm{p}}(\mathbf{r}) \approx R(\mathbf{r})/R_0$). For $\xi_{\mathrm{ch}} < \xi^s_{\mathrm{ch}} - \epsilon$, we have $H_\epsilon = 0$ and the coordinate $\xi_{\mathrm{p}}$ is equivalent to $\xi_{\mathrm{ch}}$, thus quantifying the degree of connectivity of the transmembrane defect. For more details on the definition and implementation of $\xi_{\mathrm{ch}}$ and $\xi_{\mathrm{p}}$ we refer to previous work.[18–20]

### Umbrella sampling simulations of pore nucleation and expansion

PMFs of pore formation involve a nucleation phase and an expansion phase. In this study, we carried out two sets of umbrella sampling (US) simulations using either $\xi_{\mathrm{ch}}$ or $\xi_{\mathrm{p}}$, and we combined the US windows into a single PMF. Starting points for umbrella sampling were generated using constant-velocity pulling simulations along $\xi_{\mathrm{ch}}$ from 0 to 1 over 100 ns for pore nucleation simulations and along $\xi_{\mathrm{p}}$ from 0 to 7 over 200 ns for pore expansion. A force constant of 3000 kJ/mol was used. For pore expansion simulations, flat-bottomed restraints along the membrane normal were applied to the lipid C2 atoms (in the glycerol region). The thickness of the flat-bottom region $R_{\mathrm{fb}}$ was chosen to allow for normal head group fluctuations, but to exclude large-scale membrane undulations or deformations (Table S9). The atoms used for making the membrane whole prior to computing the membrane center of mass (periodic boundary condition atoms) were chosen near the tip of the lipid tails at the center of the membrane.

For US simulations of pore nucleation along $\xi_{\mathrm{ch}}$, a spacing of 0.08 was used between US windows in the early nucleation regime ($\xi_{\mathrm{ch}} < 0.64$) with a force constant of 5000 kJ/mol and a spacing of 0.02 was used close to the nucleation barrier with a force constant of 10000 kJ/mol. For simulations of pore expansion along $\xi_\mathrm{p}$, a spacing of 0.03 was used around the pore nucleation barrier $\xi_\mathrm{p} \in (0.64, 1.12)$ with a force constant of 5000 kJ/mol, while for larger pores a spacing of 0.15 was used with a force constant of 400 kJ/mol. For pore nucleation, 27 US windows were run for 150 ns each. For pore expansion, 63 US windows were run for 50 ns for single-lipid membranes, 100 ns for binary lipid mixtures and 150 ns for complex membranes. The temperature during US simulations was kept at 310 K. For analysis of the nucleation simulations the initial 20 ns were discarded for equilibration. In case of the expansion simulations of single lipid membranes the initial 10 ns were discarded while for all membrane systems containing multiple lipid species the initial 50 ns were removed. US windows from pore nucleation in the range $\xi_\mathrm{ch} < 0.875$ were combined with the US windows of pore expansion to compute PMFs spanning both nucleation and expansion. PMFs were calculated using the GROMACS implementation of the weighted histogram method (WHAM).[23,24] Uncertainties of the PMFs were estimated by Bayesian bootstrapping of complete histograms over 50 rounds.[24]

PMF calculations along complex conformational transitions frequently suffer from poor convergence or hysteresis effects. To exclude that our PMFs are biased by such problems, we computed two PMFs both in forward or in backward direction by taking starting initial frames for umbrella sampling from constant-velocity pulling simulations of pore opening or pore closing, respectively. For two complex membranes with lipid compositions of the outer plasma membrane leaflet or of the Golgi apparatus, we find only a small difference between PMFs of pore opening and closing (Figure S1). Thus, the PMFs are reasonably converged and exhibit only minor hysteresis effects.

The free energy of pore nucleation $\Delta G_\mathrm{nuc}$ was defined as the PMF value at $\xi_\mathrm{p} = 0.92$, characterized by the presence a thin transmembrane water wire. The uncertainty of $\Delta G_\mathrm{nuc}$ was taken from the uncertainty of the PMF. The line tension $\gamma$ along the pore rim was taken from the slope of a linear fit to the PMF in the range $4 \leq \xi_\mathrm{p} \leq 6.5$. The fit was performed

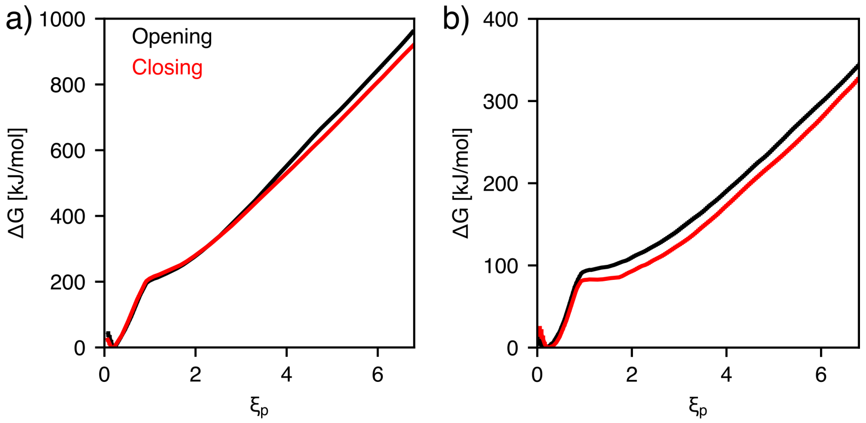


Figure S1: Comparison of PMFs along the opening (black) or closing pathway (red) for (a) membrane based on the lipid composition of the outer plasma membrane leaflet or (b) membrane of the Golgi apparatus. Opening and closing PMFs were obtained by starting umbrella sampling simulations from MD frames taken either from constant-velocity pulling simulations in opening or closing direction, respectively. Because the free energy is a state function, the PMFs should not depend on the direction of the pathway. Thus, major differences between PMFs along the opening versus the closing pathway would indicate hysteresis, which is common problem for PMF calculations along large-scale structural transitions.[25] The absence of hysteresis is a strong indication of convergence. Here, the PMFs along the opening and closing pathway are similar, suggesting that the PMFs are reasonably converged and that hysteresis is only a minor problem.

for a set of 50 bootstrapped pmf profiles and $\gamma$ and its error calculated from the average and the standard deviation.

### Additional simulation analysis

To analyze the distributions of lipids along the rim of an open pore, lipid densities were computed as function of radial distance $r$ from the pore center and of membrane normal $z$. Lipid densities were calculated from simulation restrained at $\xi_\mathrm{p} \approx 5.5$ over 400 ns for complex membranes and over 200 ns for simpler model membranes, respectively. For visualization purposes the thickness parameter $D$ used to define $\xi_\mathrm{p}$ was increased to 1.5 nm. The density distributions were obtained using an in–house modified version of the GROMACS module gmx density available at `https://gitlab.com/cbjh/gromacs-chain-coordinate`.

The thickness of the membrane core $d_\mathrm{hc}$ was derived from the mass density profiles of the hydrophobic tail atoms, which was computed from the last 50 ns of the equilibration simulations. The thickness was defined by setting a density threshold of 500 kg/m$^3$.

### Calculation of elastic properties

For the computation of elastic properties, simulations of pre-equilibrated bilayers were continued for 200 ns using Gromacs 2020.3.[8] The temperature was maintained at 303.15 K using a Nosé-Hoover thermostat,[26,27] and the pressure was maintained at 1 bar using a semi-isotropic Parrinello-Rahman barostat.[28] We used a 1.2 nm real-space cutoff for electrostatics and Lennard-Jones interactions. Long-range electrostatics were treated using the particle-mesh Ewald method.[13] Because the pressure decomposition does not support the SETTLE algorithm,[14] the triangular geometry of water was constrained using LINCS with order five.[15] No dispersion corrections or potential switching were applied. The timestep was 2 fs. 20.000 MD snapshots with velocities were taken from each simulation.

The local stress tensor was computed using the rerun functionality of the gmx mdrun module of our implementation[29] of a Goetz-Lipowsky-Decomposition[30] into the code by Sega *et al.*[31] The modified GROMACS version is available at `https://github.com/allolio`. The same cutoffs were applied as described above. Our approach is similar to the method in Ref. 32. Specifically, we set the normal pressure $p_\mathrm{N}$ to impose zero surface tension:

$$\sigma = \int_{-l}^{l} \pi(z)\,\mathrm{d}z = \int_{-l}^{l} \Big[ -\frac{1}{2}\{p_{xx}(z) + p_{yy}(z)\} + p_\mathrm{N} \Big]\,\mathrm{d}z = 0. \tag{2}$$

Here, $\sigma$ is obtained from the diagonal components of the stress tensor,[33] the integral is carried out over the entire simulation box $[-l, l]$ along the membrane normal $z$, where $z = 0$ corresponds to the center of the bilayer. Since this approach requires $p_\mathrm{N}$ being close to 1 bar, we validated that that deviations from this target were $\leq$ 1.2 bar for all simulations.

Bending modulus $\kappa$ and tilt modulus $\kappa_\theta$ were computed using the ReSIS method.[34] We used lipid director definitions and an implementation from the the LIPIDATOR-TOOLKIT available at `https://github.com/allolio/lipidator-toolkit`. This allowed the use of the well-known[35,36] relation of the first bending moment to the product of spontaneous curvature, $J_\mathrm{s}$ and $\kappa$

$$\kappa J_\mathrm{s} = \int_0^l \pi(z) z \mathrm{d}z. \tag{3}$$

Here, the integration is carried out along the membrane normal $\hat{\mathbf{e}}_z$ and only over one monolayer, and hence monolayer elastic properties are used. The reported values are averages over both monolayers. Bilayer values can be obtained by summing up the monolayer values, resulting in a bilayer $J_\mathrm{s}^\mathrm{b} = 0$ for symmetric membranes. We used spline interpolation for the integration of the pressure profiles. Lipid (mixture) data prepared specifically for this study are POPA, POPG, POPI, POPS as well as mixtures of POPC with 16-lyso-PC, oleic acid, or cholesterol. Other lipid data were taken from previous work.[29,32] New director vectors for oleic acid and lyso-PC will be added to the LIPIDATOR-TOOLKIT repository. Error bars denote standard errors computed from subsampling.

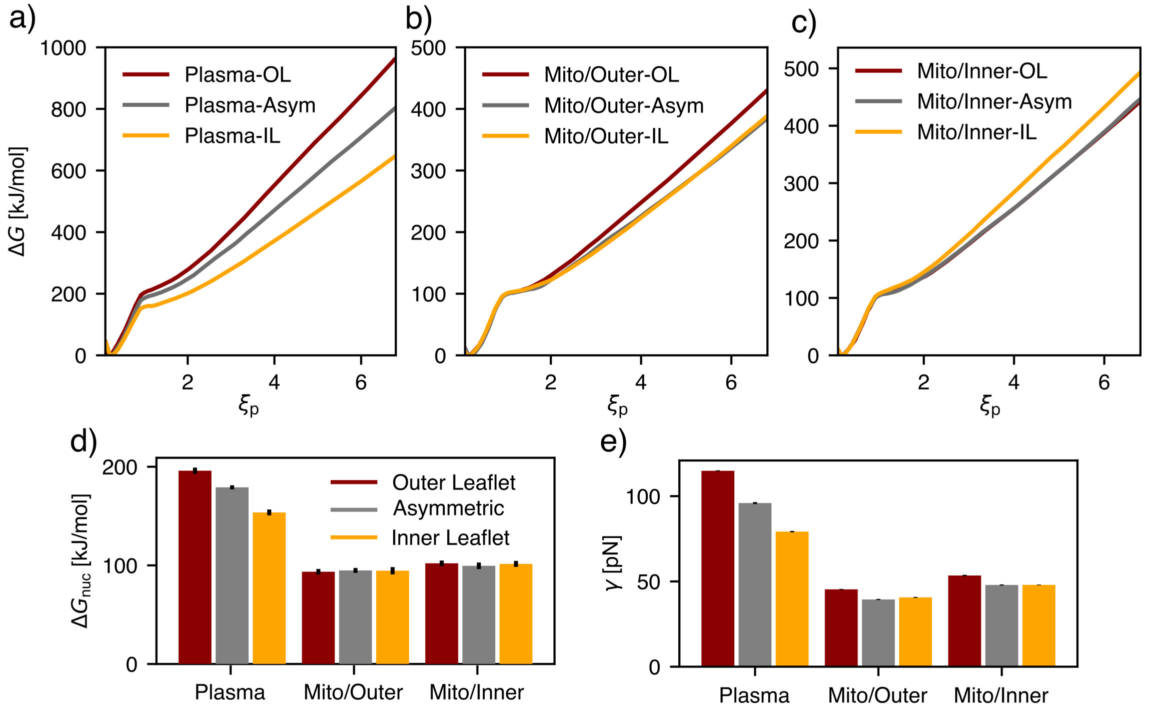


Figure S2: PMFs of pore formation for symmetric versus asymmetric membrane models. PMFs for membranes based on the composition of the outer leaflet (OL, dark red) and inner leaflet (IL, orange) are shown together with the profile of the corresponding asymmetric membranes (gray): (a) Plasma membranes, (b) outer mitochondrial membrane, and (c) inner mitochondrial membranes. (d) Bar plot of nucleation free energy $\Delta G_{\rm nuc}$ and (e) line tension $\gamma$.

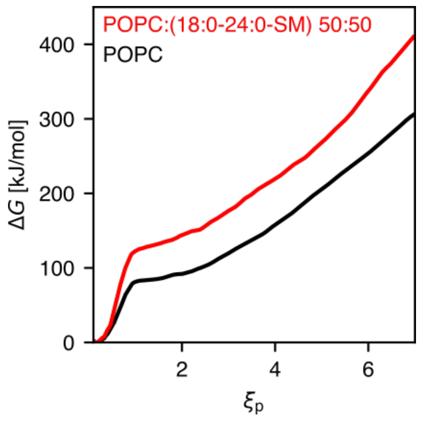


Figure S3: On the large effect of saturated sphingomyelin (18:0-24:0-SM) on pore free energy. PMF for a pure POPC membrane (black) and for a 50:50 mixture of POPC with 18:0-24:0-SM (red).

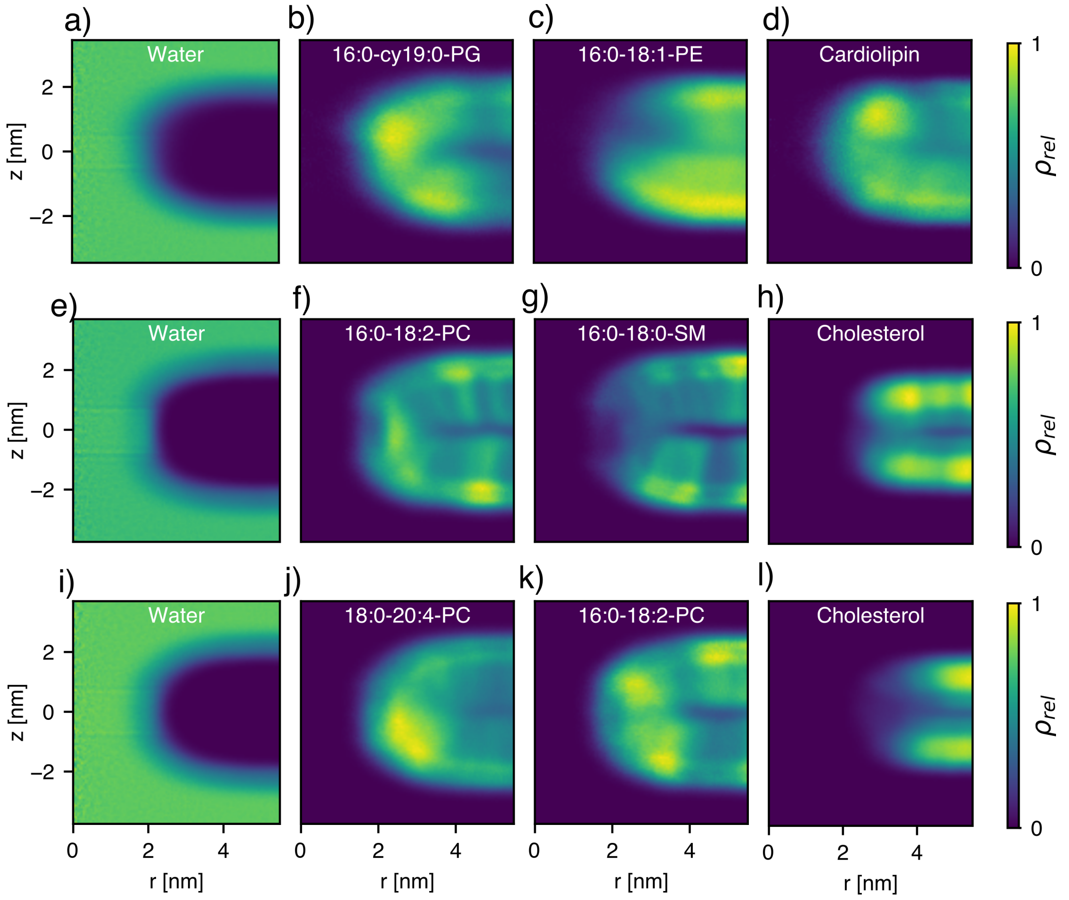


Figure S4: Relative mass densities of water and lipid species along the pore rim, plotted as function of lateral distance $r$ from the pore center and normal distance $z$ from the membrane center. Densities were averaged from extended simulations of umbrella windows with pore radii of approximately 2 nm. Densities are normalized by dividing by the maximal density of the respective constituent. Densities are shown for selected lipids (see labels) for membranes of (a-d) *E. coli* extract ($\xi_p \sim 5.7$), (e-h) outer plasma membrane leaflet ($\xi_p \sim 5.4$), and (i-l) endosome ($\xi_p \sim 5.6$). Evidently, cholesterol, sphingomyelin, and PE lipids are depleted at the pore rim (panels c, g, h, l), whereas PG lipids (panel b) and –to a lower degree– polyunsaturated PC (panels j, k) are enriched at the rim.

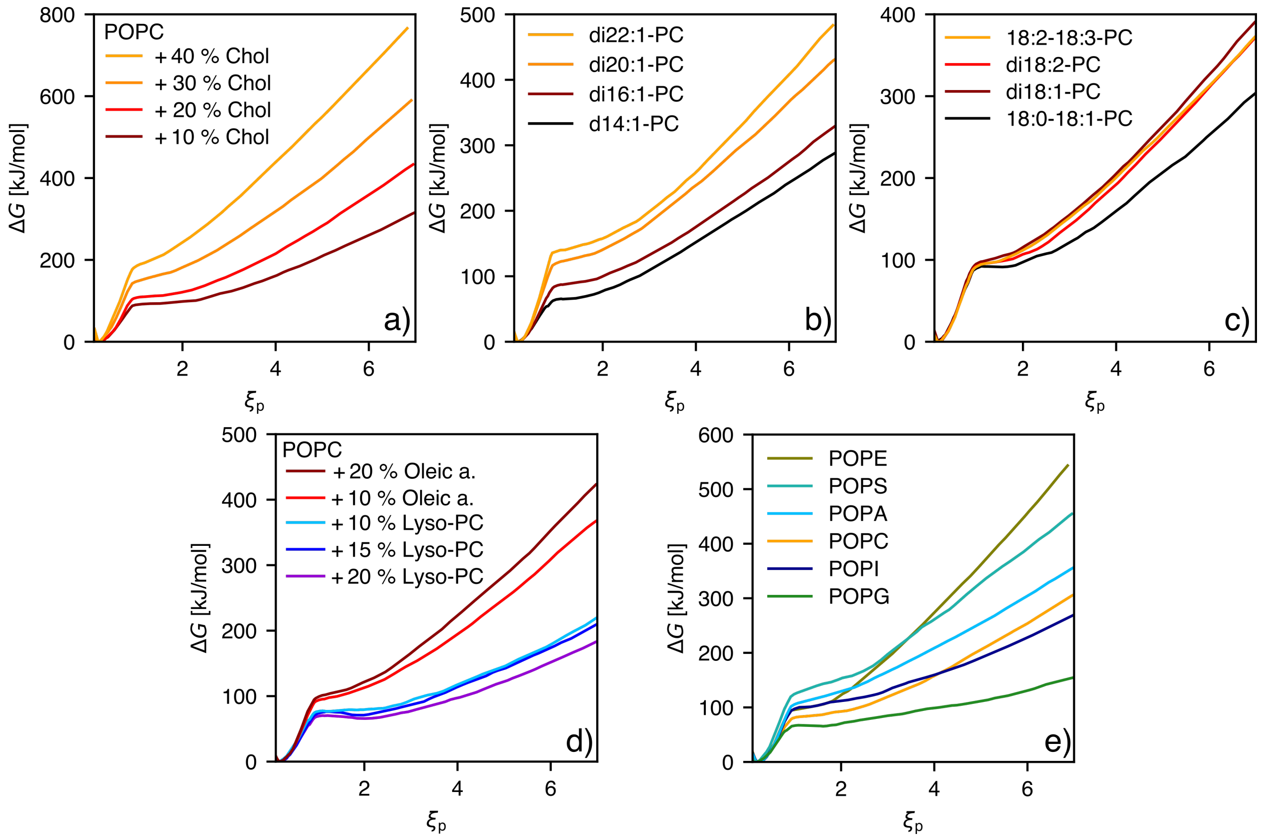


Figure S5: PMFs of model membranes with: (a) with varying sterol content, (b) with varying number of acyl tail carbons, (c) with varying number of double bonds, (d) with varying content of lyso-PC or oleic acid, (e) with varying head group composition.

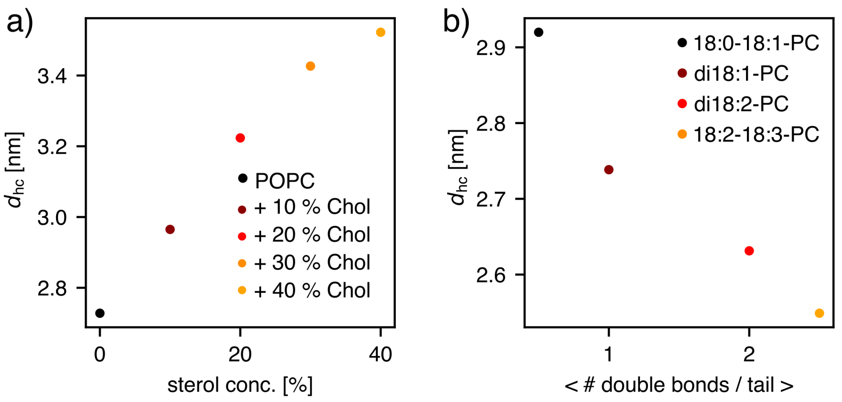


Figure S6: Effect of unsaturation and sterol content on the thickness of the membrane core $d_{\mathrm{hc}}$ of the studied model systems. (a) Effect on $d_{\mathrm{hc}}$ due to the addition of cholesterol within a POPC membrane. (b) Effect of the degree of unsaturation on $d_{\mathrm{hc}}$ using a set of PC model membranes.

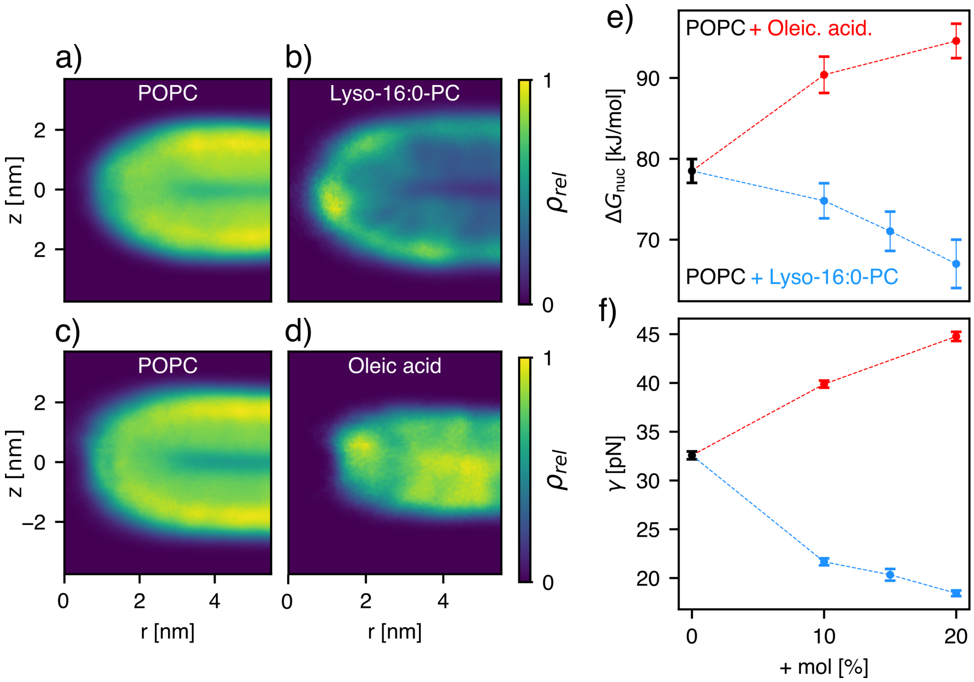


Figure S7: Influence of addition of curvature-inducing lipids. (a-d) Normalized density of lipids along the pore rim as function of radial distance $r$ from the pore center and normal distance $z$ from the membrane center for (a/b) simulation of POPC–lyso-16:0-PC 80:20 and (c/d) POPC–oleic acid 80:20. Lyso-PC and oleic acid are enriched or depleted at the pore rim, respectively. (e) Effect of increasing oleic acid or lyso-PC concentration on $\Delta G_{\mathrm{nuc}}$ and (f) $\gamma$.

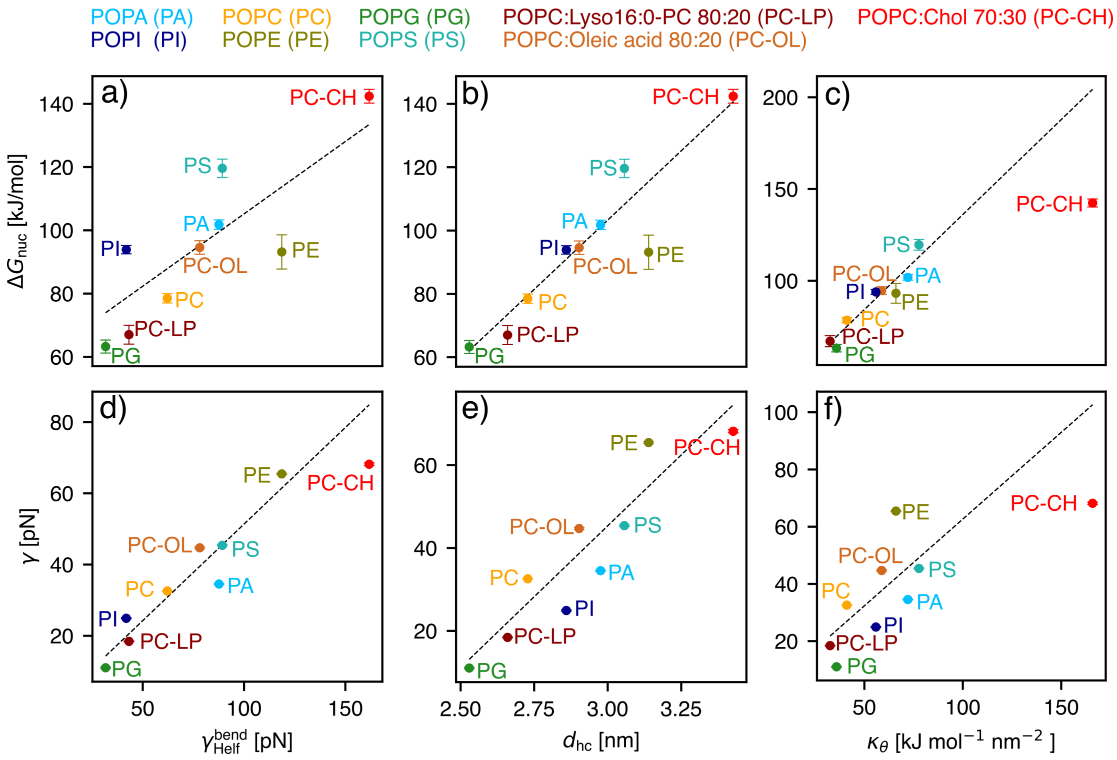


Figure S8: Correlations of $\Delta G_{\mathrm{nuc}}$ or $\gamma$ with $\gamma_{\mathrm{Helf}}^{\mathrm{bend}}$, hydrophobic core thickness $d_{\mathrm{hc}}$ and tilt modulo $\kappa_0$ (see axis labels) for model membranes (see Figure 6 in main text) including mixture of 70 % POPC with 30 % cholesterol (red). Results of linear regressions are indicated as dashed lines. Note that the data for the POPC/cholesterol mixture was not included in the regression model.

Table S1: Lipid composition of plasma membrane systems of small ($N_{\text{small}}$) and large ($N_{\text{large}}$) membrane systems of the asymmetric membranes for the outer leaflet (OL) and inner leaflet (IL) and for the symmetric membrane based on the outer leaflet (OL-sym).

| Component | OL | | IL | | OL-sym | |
|---|---|---|---|---|---|---|
| | $N_{\text{small}}$ | $N_{\text{large}}$ | $N_{\text{small}}$ | $N_{\text{large}}$ | $N_{\text{small}}$ | $N_{\text{large}}$ |
| 16:0-16:0-PC (DPPC) | - | - | 4 | 6 | - | - |
| 16:0-18:1-PC (POPC) | 9 | 16 | 5 | 9 | 8 | 15 |
| 16:0-18:2-PC (PLPC) | 25 | 48 | 16 | 30 | 24 | 45 |
| 16:0-22:4-PC (PAPC) | 9 | 16 | - | - | 8 | 15 |
| 18:1-16:0-SM (PSM) | 20 | 38 | 4 | 6 | 20 | 36 |
| 18:1-24:1-SM (NSM) | 11 | 21 | - | - | 12 | 21 |
| 18:1-24:0-SM (LSM) | 11 | 21 | - | - | 12 | 21 |
| 18:0-22:4-PI (SAPI) | 3 | 5 | 4 | 6 | 4 | 6 |
| 16:0-22:4-PS (PAPS) | 6 | 11 | 39 | 74 | 6 | 9 |
| 16:0-22:4-PE (PAPE) | 3 | 5 | - | - | 4 | 6 |
| 16:0-18:1-PE (POPE) | - | - | 2 | 3 | - | - |
| 16:0-22:6-PE (PDoPE) | - | - | 16 | 30 | - | - |
| 18:1-22:4-PE (OAPE) | - | - | 5 | 9 | - | - |
| Cholesterol | 73 | 136 | 63 | 124 | 64 | 126 |
| Sum | 170 | 317 | 160 | 300 | 162 | 300 |

Table S2: Lipid composition of endosomal membrane of the small ($N_{\text{small}}$) and large ($N_{\text{large}}$) simulation systems.

| Component | $N_{\text{small}}$ | $N_{\text{large}}$ |
|---|---|---|
| 16:0-18:2-PC (PLPC) | 8 | 30 |
| 18:0-22:4-PC (SAPC) | 14 | 51 |
| 18:0-22:6-PC (SDPC) | 5 | 18 |
| 16:0-18:2-PE (PLPE) | 2 | 6 |
| 16:0-22:6-PE (PDoPE) | 4 | 12 |
| 18:0-22:4-PE (SAPE) | 6 | 21 |
| 18:0-22:4-PI (SAPI) | 5 | 18 |
| 18:0-22:6-PI (SDPI) | 2 | 6 |
| 18:0-24:0-SM (DSM) | 7 | 24 |
| 18:1-18:1-SM (OSM) | 7 | 24 |
| Cholesterol | 21 | 90 |
| Sum | 81 | 300 |

Table S3: Lipid composition of lysosomal membranes of the small ($N_{\text{small}}$) and large ($N_{\text{large}}$) simulation systems.

| Component | $N_{\text{small}}$ | $N_{\text{large}}$ |
|---|---|---|
| 16:0-18:2-PC (PLPC) | 11 | 33 |
| 18:0-22:4-PC (SAPC) | 18 | 53 |
| 18:0-22:6-PC (SDPC) | 6 | 18 |
| 16:0-18:2-PE (PLPE) | 7 | 21 |
| 18:0-22:4-PE (SAPE) | 18 | 53 |
| 18:0-22:4-PI (SAPI) | 6 | 18 |
| 18:0-22:6-PI (SDPI) | 2 | 6 |
| 18:0-22:4-PS (SAPS) | 2 | 6 |
| 18:0-22:6-PS (SDPS) | 1 | 3 |
| 18:1-16:0-SM (PSM) | 3 | 9 |
| 18:1-24:0-SM (LSM) | 3 | 9 |
| 18:1-18:1-BMP (BMGP) | 7 | 17 |
| Cholesterol | 18 | 53 |
| Sum | 102 | 300 |

Table S4: Lipid composition of ER membranes of the small ($N_{\text{small}}$) and large ($N_{\text{large}}$) simulation systems.

| Component | $N_{\text{small}}$ | $N_{\text{large}}$ |
|---|---|---|
| 16:0-18:1-DAG (POGL) | 4 | 12 |
| 18:1-18:1-DAG (DOGL) | 3 | 9 |
| 16:1-16:1-PE (DYPE) | 17 | 51 |
| 16:1-16:1-PC (DYPC) | 20 | 60 |
| 16:0-18:1-PC (POPC) | 5 | 15 |
| 16:0-18:1-PI (POPI) | 25 | 75 |
| 16:0-18:1-PA (POPA) | 2 | 6 |
| 18:1-18:1-PA (DOPA) | 1 | 3 |
| 16:0-18:1-PS (POPS) | 2 | 6 |
| 18:1-18:1-PS (DOPS) | 1 | 3 |
| Ergosterol | 10 | 30 |
| Sum | 100 | 300 |

Table S5: Lipid composition of Golgi membranes of the small ($N_{\mathrm{small}}$) and large ($N_{\mathrm{large}}$) simulation systems.

| Component | $N_{\mathrm{small}}$ | $N_{\mathrm{large}}$ |
|---|---|---|
| 16:0-18:1-PC (POPC) | 11 | 33 |
| 16:0-18:2-PC (PLPC) | 14 | 42 |
| 18:0-22:4-PC (SAPC) | 20 | 60 |
| 16:0-18:2-PE (PLPE) | 4 | 12 |
| 16:0-18:0-PE (PSPE) | 8 | 24 |
| 18:0-22:4-PE (SAPE) | 5 | 15 |
| 16:0-18:0-PI (PSPI) | 2 | 6 |
| 16:0-18:1-PI (POPI) | 5 | 15 |
| 18:0-22:4-PI (SAPI) | 2 | 6 |
| 16:0-18:1-PS (POPS) | 4 | 12 |
| 18:0-22:0 -SM (TSM) | 12 | 36 |
| Lyso-16:0-PC | 5 | 15 |
| Cholesterol | 8 | 24 |
| Sum | 100 | 300 |

Table S6: Lipid composition of outer mitochondrial membranes for outer leaflet (OL) and inner leaflet (IL), each of the small ($N_{\mathrm{small}}$) and large ($N_{\mathrm{large}}$) simulation systems.

| Component | OL | | IL | |
|---|---|---|---|---|
| | $N_{\mathrm{small}}$ | $N_{\mathrm{large}}$ | $N_{\mathrm{small}}$ | $N_{\mathrm{large}}$ |
| 16:0-18:1-PC (POPC) | 6 | 18 | 6 | 18 |
| 16:0-18:2-PC (PLPC) | 18 | 54 | 18 | 54 |
| 18:0-22:4-PC (SAPC) | 30 | 90 | 30 | 90 |
| 16:0-18:1-PE (POPE) | 4 | 12 | 2 | 6 |
| 16:0-18:2-PE (PLPE) | 9 | 27 | 5 | 15 |
| 18:0-22:4-PE (SAPE) | 25 | 75 | 13 | 39 |
| 18:0-22:4-PI (SAPI) | 4 | 12 | 22 | 66 |
| 18:0-22:4-PS (SAPS) | 1 | 3 | 3 | 9 |
| 16:0-18:1-PA (POPA) | 1 | 3 | 1 | 3 |
| 18:2-18:2/18:2-18:2/CL (TLCL1) | 2 | 4 | - | - |
| Sum | 100 | 300 | 100 | 300 |

Table S7: Lipid composition of inner mitochondrial membranes for outer and inner leaflets.

| Component | OL | | IL | |
|---|---|---|---|---|
| | $N_{\text{small}}$ | $N_{\text{large}}$ | $N_{\text{small}}$ | $N_{\text{large}}$ |
| 16:0-18:1-PC (POPC) | 7 | 21 | 3 | 9 |
| 16:0-18:2-PC (PLPC) | 19 | 57 | 10 | 30 |
| 18:0-22:4-PC (SAPC) | 32 | 96 | 16 | 48 |
| 16:0-18:1-PE (POPE) | 5 | 15 | 4 | 12 |
| 16:0-18:2-PE (PLPE) | 8 | 24 | 8 | 24 |
| 18:0-22:4-PE (SAPE) | 24 | 72 | 24 | 72 |
| 18:0-22:4-PE (SAPI) | 5 | 15 | 6 | 18 |
| 18:0-22:4-PE (SAPS) | 3 | 9 | 3 | 9 |
| 18:2-18:2/18:2-18:2/CL (TLCL1) | 11 | 33 | 26 | 78 |
| Sum | 114 | 342 | 100 | 300 |

Table S8: Lipid composition of *E. coli* polar lipid extract of the small ($N_{\text{small}}$) and large ($N_{\text{large}}$) simulation systems.

| Component | $N_{\text{small}}$ | $N_{\text{large}}$ |
|---|---|---|
| 16:1-16:0/cy17:0-cy19:0-CL (YPMNCL) | 1 | 3 |
| 16:1-16:0/cy17:0-18:1-CL (YPMVCL) | 1 | 3 |
| cy17:0-16:0/16:0-18:1-CL (MPPVCL) | 2 | 6 |
| 17:0-16:0/16:0-17:0-CL (MPPMCL) | 1 | 3 |
| 16:0-cy19:0-PG (PMPG) | 10 | 36 |
| 16:0-cy19:0-PG (PNPG) | 2 | 6 |
| 16:1-18:1-PG YOPG | 1 | 3 |
| 16:0-18:1-PG (POPG) | 4 | 12 |
| 16:0-16:1-PG (PYPG) | 2 | 6 |
| 16:1-18:1-PE (YOPE) | 12 | 45 |
| 16:0-cy17:0-PE (PMPE) | 15 | 54 |
| 16:0-cy19:0-PE (PNPE) | 5 | 18 |
| cy17:0-18:1-PE (MVPE) | 7 | 27 |
| 16:0-18:1-PE (POPE) | 17 | 77 |
| Sum | 81 | 300 |

Table S9: Parameters of pore reaction coordinate for different systems. Number of slices of transmembrane cylinder used to define $\xi_{\mathrm{ch}}$ during umbrella sampling of pore nucleation $N_{\mathrm{s}}^{\mathrm{nuc}}$ and during pore expansion $N_{\mathrm{s}}^{\mathrm{exp}}$, radius of the flat-bottomed region $R_{\mathrm{fb}}$ of the flat-bottomed potential used to exclude large-scale membrane undulations, and radius $R_0$ of a thin nucleated pore used to define $\xi_{\mathrm{p}}$. $R_0$ is not a free parameter but set automatically at the beginning of the simulation based on the number of slices ($N_{\mathrm{s}}^{\mathrm{nuc}}$ or $N_{\mathrm{s}}^{\mathrm{exp}}$) and $D$, see Ref. 18.

| System | $N_{\mathrm{s}}^{\mathrm{nuc}}$ | $N_{\mathrm{s}}^{\mathrm{exp}}$ | $R_{\mathrm{fb}}$ [nm] | $R_0$ [nm] |
|---|---|---|---|---|
| Plasma-Asym | 44 | 44 | 2.1 | 0.326115 |
| Plasma-OL | 44 | 45 | 2.2 | 0.332651 |
| Plasma-IL | 43 | 44 | 2.1 | 0.326115 |
| E. coli extract | 35 | 34 | 2.0 | 0.372404 |
| Endosome | 41 | 43 | 2.1 | 0.335857 |
| ER | 29 | 37 | 2.0 | 0.358956 |
| Lysosome | 37 | 38 | 2.0 | 0.353815 |
| Golgi | 34 | 36 | 2.0 | 0.363457 |
| Mito/Outer-OL | 31 | 32 | 2.0 | 0.380517 |
| Mito/Outer-IL | 31 | 31 | 2.0 | 0.383527 |
| Mito/Outer-Asym | 33 | 33 | 2.0 | 0.374414 |
| Mito/Inner-OL | 32 | 35 | 2.0 | 0.365406 |
| Mito/Inner-IL | 33 | 35 | 2.0 | 0.365406 |
| Mito/Inner-Asym | 33 | 34 | 2.0 | 0.372404 |
| (16:0-18:1)-PC (POPC) | 25 | 27 | 2.0 | 0.39796 |
| (16:0-18:1)-PA (POPA) | 32 | 33 | 2.0 | 0.374414 |
| (16:0-18:1)-PE (POPE) | 34 | 34 | 2.3 | 0.372404 |
| (16:0-18:1)-PI (POPI) | 30 | 32 | 2.0 | 0.380517 |
| (16:0-18:1)-PG (POPG) | 23 | 22 | 2.0 | 0.426354 |
| (16:0-18:1)-PS (POPS) | 32 | 32 | 2.3 | 0.380517 |
| POPC:Chol 90:10 | 27 | 32 | 2.3 | 0.380517 |
| POPC:Chol 80:20 | 34 | 31 | 2.3 | 0.383527 |
| POPC:Chol 70:30 | 37 | 37 | 2.3 | 0.358956 |
| POPC:Chol 60:40 | 40 | 42 | 2.3 | 0.344646 |
| POPC:Lyso-16:0-PC 80:20 | 22 | 25 | 2.0 | 0.406944 |
| POPC:Lyso-16:0-PC 85:15 | 22 | 28 | 2.0 | 0.396042 |
| POPC:Lyso-16:0-PC 90:10 | 23 | 26 | 2.0 | 0.38848 |
| POPC:Oleic acid 80:20 | 30 | 31 | 2.3 | 0.38848 |
| POPC:Oleic acid 90:10 | 28 | 28 | 2.3 | 0.38848 |
| POPC:(18:0-24:0)-SM 50:50 | 27 | 30 | 2.3 | 0.38848 |
| 14:1-14:1-PC (DRPC) | 19 | 19 | 1.7 | 0.438885 |
| 16:1-16:1-PC (DYPC) | 24 | 27 | 1.9 | 0.39796 |
| 18:1-18:1-PC (DOPC) | 28 | 29 | 2.0 | 0.390468 |
| 20:1-20:1-PC (DGPC) | 32 | 35 | 2.2 | 0.365406 |
| 22:1-22:1-PC (DEPC) | 38 | 39 | 2.6 | 0.354824 |
| 12:0-12:0-PC (DLPC) | 14 | 16 | 1.7 | 0.461476 |
| 18:0-18:1-PC (SOPC) | 25 | 32 | 2.0 | 0.380517 |
| 18:2-18:2-PC (DUPC) | 26 | 29 | 2.0 | 0.390468 |
| 18:2-18:3-PC (LLPC) | 26 | 28 | 2.0 | 0.396042 |

Table S10: Results of linear regression $Y = m \cdot X + n$ to estimate correlation between $\Delta G_{\text{nuc}}$ and $\gamma$ with elastic/geometric properties.

| $Y$ | $X$ | $m$ | $n$ | R |
|---|---|---|---|---|
| $\Delta G_{\text{nuc}}$ | $\gamma_{\text{Helf}}^{\text{bend}}$ | 0.5 ± 0.2 | 59 ± 14 | 0.680 |
| $\gamma$ | $\gamma_{\text{Helf}}^{\text{bend}}$ | 0.54 ± 0.08 | -3 ± 5 | 0.947 |
| $\Delta G_{\text{nuc}}$ | $d_{\text{hc}}$ | 88 ± 15 | -159 ± 43 | 0.920 |
| $\gamma$ | $d_{\text{hc}}$ | 68 ± 16 | -160 ± 45 | 0.867 |
| $\Delta G_{\text{nuc}}$ | $\kappa_\theta$ | 1.04 ± 0.13 | 33 ± 7 | 0.958 |
| $\gamma$ | $\kappa_\theta$ | 0.61 ± 0.26 | 2.024 ± 14 | 0.687 |

Table S11: Experimental measurements of line tension $\gamma$ reported previously.

| membrane composition | $\gamma$ [pN] | Reference |
|---|---|---|
| Egg PC | 14.2 ± 0.7 | Portet T. & Dimova R.A. 2010[37] |
| Egg PC | 8.6 ± 0.4 | Chernomordik L. *et al.* 1985[38] |
| DOPC | 28 ± 3 | Portet T. & Dimova R.A. 2010[37]sun2022physical |
| DOPC from Avanti | 20 ± 4 | Karatekin E. *et al.* 2003[39] |
| DOPC from Sigma | 6.9 ± 0.4 | Karatekin E. *et al.* 2003[39] |
| E.coli PE | 16.0 ± 0.6 | Chernomordik L. *et al.* 1985[38] |
| SOPC | 9.2 ± 0.7 | Zhelev D.V. *et al.* 1993[40] |
| DOPG/DOPE (4/6) | 15.6 ± 0.7 | Tazawa K. & Yamazaki M. 2023[41] |
| DOPG/DOPC (4/6) | 10.7 ± 0.4 | Tazawa K. & Yamazaki M. 2023[41] |
| DOPG/DOPC/LPC (40/55/10) | 6.7 ± 0.2 | Tazawa K. & Yamazaki M. 2023[41] |
| POPC | 40 ± 6 | Lira R.B. *et al.* 2021[42] |
| POPC/POPG (1/1) | 23 ± 6 | Lira R.B. *et al.* 2021[42] |

Table S12: Pore nucleation free energies and line tension for all simulated membranes.

| System | $\Delta G_{\mathrm{nuc}}$ [kJ/mol] | $\gamma$ [pN] |
|---|---|---|
| Plasma-Asym | 179 ± 2 | 96.0 ± 0.6 |
| Plasma-OL | 196 ± 3 | 114.9 ± 0.3 |
| Plasma-IL | 154 ± 3 | 79.2 ± 0.4 |
| E. coli extract | 105 ± 4 | 51.38 ± 0.22 |
| Endosome | 138 ± 5 | 69.7 ± 0.4 |
| ER | 106 ± 2 | 50.8 ± 0.5 |
| Lysosome | 109 ± 3 | 48.3 ± 0.4 |
| Golgi | 90 ± 2 | 39.57 ± 0.16 |
| Mito/Outer-OL | 94 ± 4 | 40.64 ± 0.28 |
| Mito/Outer-IL | 94 ± 3 | 45.3 ± 0.24 |
| Mito/Outer-Asym | 95 ± 3 | 39.4 ± 0.4 |
| Mito/Inner-OL | 102 ± 3 | 53.5 ± 0.5 |
| Mito/Inner-IL | 102 ± 4 | 44.98 ± 0.19 |
| Mito/Inner-Asym | 100 ± 4 | 48.0 ± 0.4 |
| 16:0-18:1-PC (POPC) | 79 ± 2 | 32.6 ± 0.4 |
| 16:0-18:1-PA (POPA) | 102 ± 2 | 34.6 ± 0.6 |
| 16:0-18:1-PE (POPE) | 93 ± 6 | 65.5 ± 0.7 |
| 16:0-18:1-PI (POPI) | 94 ± 2 | 25.0 ± 0.4 |
| 16:0-18:1-PG (POPG) | 63 ± 3 | 11.1 ± 0.5 |
| 16:0-18:1-PS (POPS) | 120 ± 3 | 45.4 ± 0.5 |
| POPC:Chol 90:10 | 88 ± 2 | 35.0 ± 0.5 |
| POPC:Chol 80:20 | 104 ± 2 | 50.4 ± 0.7 |
| POPC:Chol 70:30 | 142 ± 3 | 68.3 ± 0.8 |
| POPC:Chol 60:40 | 177 ± 4 | 88.4 ± 0.6 |
| POPC:Lyso-16:0-PC 80:20 | 67 ± 3 | 18.4 ± 0.3 |
| POPC:Lyso-16:0-PC 85:15 | 71 ± 3 | 20.3 ± 0.6 |
| POPC:Lyso-16:0-PC 90:10 | 75 ± 3 | 21.7 ± 0.4 |
| POPC:Oleic acid 80:20 | 95 ± 3 | 44.8 ± 0.5 |
| POPC:Oleic acid 90:10 | 90 ± 3 | 39.9 ± 0.4 |
| POPC:(18:0-24:0)-SM | 119 ± 4 | 43.0 ± 0.7 |
| 14:1-14:1-PC (DRPC) | 63 ± 5 | 27.4 ± 0.6 |
| 16:1-16:1-PC (DYPC) | 82 ± 4 | 33.6 ± 0.4 |
| 18:1-18:1-PC (DOPC) | 91 ± 3 | 41.8 ± 0.6 |
| 20:1-20:1-PC (DGPC) | 117 ± 4 | 46.8 ± 0.4 |
| 22:1-22:1-PC (DEPC) | 136 ± 3 | 56.4 ± 0.6 |
| 12:0-12:0-PC (DLPC) | 39 ± 4 | 15.3 ± 0.6 |
| 18:0-18:1-PC (SOPC) | 87 ± 3 | 32.1 ± 0.4 |
| 18:2-18:2-PC (DUPC) | 88 ± 3 | 40.5 ± 0.3 |
| 18:2-18:3-PC (LLPC) | 88 ± 4 | 37.62 ± 0.23 |
